# MetaGEAR Explorer: rapid interactive searches and cross-cohort microbiome analyses to identify disease associations of microbial genes

**DOI:** 10.64898/2026.03.30.715271

**Authors:** Emilio Ríos, Shen Jin, Chunyang Zhang, Florian Neuhaus, Xiheng He, Svenja Weißenberger, Melanie Schirmer

## Abstract

**Background:** Investigating functional gut microbiome signals remains challenging due to the large scale and sparsity of gene-level metagenomic profiles and reproducibility of microbial gene associations across microbiome studies is limited. Further, interactive tools for phenotype-aware cross-study meta-analysis are currently lacking.

**Results:** We developed MetaGEAR Explorer, a web platform for interactive and programmatic gene-centric analyses that enables users to rapidly search genes of interest across 33 million gene families from 24 metagenomic cohorts spanning inflammatory bowel disease (IBD), colorectal cancer (CRC), and healthy individuals. To demonstrate our platform’s capabilities, we first used *narG*, a well-characterized nitrate reductase gene in Enterobacteriaceae (including *Escherichia coli* and *Klebsiella* species), to evaluate the detection of remotely related, disease-relevant homologs. While sequence-based searches using the *E. coli narG* gene as a query failed to capture the known diversity of *narG*, MetaGEAR Explorer’s domain-based search expansion identified 160 NarG-like gene families with consistent IBD enrichment across cohorts. This included *narG* homologs from *Veillonella parvula* and *Veillonella atypica* that have been recently implicated in intestinal inflammation. We further applied MetaGEAR Explorer to investigate the under-explored diversity of *clbS*, the self-protection gene against the bacterial genotoxin colibactin, which is implicated in CRC tumorigenesis. The canonical *clbS*is part of the *E. coli clb* operon, which produces colibactin. Our analysis showed that *E. coli* -associated *clbS* was rare in healthy individuals but increased in disease (1.85% in Healthy, 6.21% in IBD, and 7.62% in CRC). In contrast, domain-based expansion revealed widespread *clbS* -domain (DUF1706) homologs, present in 95.15% of healthy individuals, while disease-associated contributors shift from commensal Clostridia to Gammaproteobacteria. Notably, *Proteus mirabilis* was identified as a novel potential ClbS-like carrier enriched in IBD. Furthermore, much of this prevalence was driven by a single unclassified gene family, present in 63.40% of healthy individuals.

**Conclusions:** MetaGEAR Explorer facilitates rapid cross-cohort functional meta-analyses and can identify reproducible, biologically interpretable microbial gene signatures in IBD and CRC, such as newly identified sequence–domain dynamics in nitrate respiration and colibactin self-protection.

## 1 Background

The human gut microbiome plays a central role in health and disease [1], and perturbations in this microbial ecosystem have been implicated in disease, including inflammatory bowel disease (IBD) and colorectal cancer (CRC) [2–5]. Shotgun metagenomic sequencing data enables gene-level functional and strain-level taxonomic profiling across large patient cohorts [6, 7], yet comparing results across studies remains challenging: inconsistent taxonomic annotations across profiling methods and the extreme scale and sparsity of gene-level data (often tens of millions of genes across hundreds of samples) limit the identification of robust, reproducible microbial gene–disease associations.

Reference-based functional profilers such as HUMAnN 3 [8] quantify gene-family and pathway abundances by mapping metagenomic reads to curated reference resources (e.g., UniRef gene families and MetaCyc metabolic pathways), while sketch-based approaches such as fmh-funprofiler [9] rapidly detect reference ortholog groups (e.g., KEGG KOs) using FracMinHash-style workflows. Furthermore, curatedMetagenomicData v3 (cMD3) complements these methods by providing uniformly processed MetaPhlAn 3 and HUMAnN 3 profiles together with manually curated sample metadata across *>*22k public human microbiome metagenomes, enabling cross-cohort and phenotype-aware analyses without the need to reprocess raw sequencing data [10, 11]. These approaches are fundamentally reference-driven, and sequences that do not have a database match cannot be functionally characterized and typically remain unassigned. To capture microbial diversity absent from standard reference databases, large resources of de novo assembled sequences, such as the Integrated Gene Catalog (IGC; 9,879,896 non-redundant gut microbial genes from 1,267 metagenomes) [12] and the Unified Human Gastrointestinal Genome (UHGG) and Unified Human Gastrointestinal Protein (UHGP) collections (*>*170M gut proteins; 20.2M protein clusters at 95% identity) [13], have vastly expanded the known sequence space. However, navigation and rapid extraction of biological insights from these massive datasets remains a major challenge. While some platforms provide basic interactive access (e.g., MetaGeneBank provides a disease-filtered web interface with gene-sequence search and abundance summaries over 4,470 gut metagenomes [14]), they lack deeper multi-study and genomic integration. An integrated system that combines interactive, phenotype-stratified exploration of microbial genes with catalog-scale contextualization (e.g., genomic neighborhood and pangenome views) across multiple cohorts is still missing (Table 1).

**Table 1.** Comparison of MetaGEAR Explorer with representative resources for functional metagenomic analyses. The first row labels each tool by data-generation strategy (**Ref.** = reference-based; **De novo** = de novo assembly-based). curatedMetagenomic-Data v3 (cMD3) distributes uniformly processed MetaPhlAn 3 and HUMAnN 3 profiles with manually curated sample metadata across >22k metagenomes. IGC and UHGG/UHGP are de novo gene/protein catalogs distributed primarily for bulk download; some public portals provide online access (e.g., CNGBdb for IGC; MGnify for UHGG/UHGP-related catalogs).

| Feature | cMD3 | IGC | UHGP | MetaGEAR Explorer |
| --- | --- | --- | --- | --- |
| Strategy | Ref. | De novo | De novo | De novo |
| Analysis resolution | Species, gene families & pathways | Gene clusters | Protein sequences/clusters | Gene & protein families |
| Built-in cross-cohort dataset integration | ✓ | – | – | ✓ |
| Disease/phenotype stratification supported | ✓ | – | – | ✓ |
| Interactive web interface | – | ✓<sup>†</sup> | ✓<sup>‡</sup> | ✓ |
| Sequence-based query (e.g., BLAST/HMMER) | – | ✓<sup>†</sup> | ✓<sup>‡</sup> | ✓ |
| Domain/HMM-based search | – | – | – | ✓ |
| Genomic neighborhood context linked to hits | – | – | ✓<sup>‡</sup> | ✓ |
| Cross-cohort evidence summary | – | – | – | ✓ |
| Programmatic access | R/Bioconductor + CLI | FTP/HTTP | FTP | REST |
| Scale (integrated samples/genomes) | 22 588* | 1 267 | – | 9 053 |
| Scale (de novo catalog size) | – | 9.88M genes | 170.6M proteins (UHGP-100) | 33M genes / 23M proteins |

\*cMD3 is reported as 22 588 samples in the package and 22 710 samples analyzed in the companion resource paper.
<sup>†</sup> CNGBdb hosts IGC downloads and provides a web BLAST interface and abundance/profile tables.
<sup>‡</sup>UHGP is distributed via MGnify alongside the companion UHGG genome catalog; MGnify provides web browsing/search tools for genome catalogs and large protein resources.

Here, we introduce MetaGEAR Explorer, an easily accessible platform that eliminates the need for local computational resources by allowing researchers to explore microbial gene functions and disease associations across a comprehensive collection of IBD, CRC and healthy metagenomic cohorts within seconds. IBD and CRC are two major gastrointestinal diseases characterized by microbiome perturbations. While emerging evidence demonstrates causal links between specific microbial alterations and disease pathogenesis, such as the role of microbial genotoxins in CRC tumorigenesis, the underlying host-microbial mechanisms still remain largely elusive. Uncovering these mechanisms requires high-resolution functional microbiome profiles that go beyond taxonomic associations to identify the specific microbial genes that may contribute to disease development. MetaGEAR Explorer integrates uniformly processed data from 9,053 samples across 24 healthy and disease cohorts into a searchable database of 33 million gene families and 23 million protein families. Researchers can query this database by nucleotide or amino acid sequences (BLAST) or by Pfam domain configurations, and instantly explore the results through disease-stratified prevalence profiles, cross-cohort association summaries, species-level taxonomy, genomic neighborhood context, and metagenomic species pangenome (MSP) assignments. Users can transition from sequence-based searches to domain-based searches using the Pfam domains detected in the matched gene families, bridging the two search modes to uncover close as well as distant functional homologs. Furthermore, a dedicated MSP catalog browser enables organism-level exploration. All platform features are accessible through an interactive web interface and programmatic web services (RESTful API) that enables automated data retrieval and integration into custom analytical pipelines.

We demonstrate MetaGEAR Explorer’s capabilities through an analysis of the nitrate reductase-encoding *narG* gene, where a single sequence query expanded to 160 domain-matched gene families with reproducible IBD enrichment across independent cohorts. In a complementary case study of the colibactin self-protection gene *clbS*, aggregating multiple sequence variants revealed a two-layered disease pattern. *Escherichia coli* - encoded *clbS* expanded in IBD and CRC, whereas the broader DUF1706 domain family, carried largely by commensal Clostridia and a dominant unclassified family in healthy individuals, was depleted in IBD, generating new insight into sequence-to-domain decoupling at the gene-family level.

## 2 Methods

### 2.1 A uniformly processed multi-cohort gene, protein, and pangenome catalog

MetaGEAR Explorer leverages a comprehensive multi-cohort database (previously described in [15]), offering search capabilities through a web interface to investigate genes and proteins of interest, including their disease relevance. Briefly, data from 9,053 metagenomic samples spanning 24 cohorts (Supplementary Table 1) were uniformly processed through an integrated assembly- and reference-based pipeline to systematically predict, cluster, and annotate de novo assembled genes, resulting in comprehensive, non-redundant catalogs with unified taxonomic and functional profiles. The assembly-based analysis resulted in a multi-cohort database structured into a multi-cohort gene catalog (MCGC), a multi-cohort protein catalog (MCPC), and a multi-cohort metadata database. The MCGC includes 33,353,858 non-redundant gene families clustered at 95% sequence similarity and 90% coverage, and the MCPC includes 23,475,122 non-redundant protein families clustered at 90% sequence similarity and 80% coverage. Pfam domain architectures were annotated for the representative protein sequences using InterProScan 5.47-82.0. Gene families were further organized into 13,795 metagenomic species pangenomes (MSPs) using co-abundance binning with MSPminer 1.1.3 [16], with each gene classified as core, accessory, or singleton based on its presence pattern across samples within a cohort. Per-cohort sample inclusion was determined by the availability of raw sequencing data, complete metadata, and successful processing through all pipeline stages.

Of the 9,053 metagenomes, 8,744 were assigned to one of three disease groups (Healthy: 5,730; IBD: 2,384; CRC: 630) and used in the cross-cohort prevalence and disease-association analyses reported throughout this work; the remaining 309 samples carried other phenotypes or had partially incomplete metadata and contribute to the gene and protein catalogs but were excluded from the group-level disease summaries.

To enable efficient data storage and retrieval, we stored these gene and protein catalogs together with information about their abundance, taxonomy, function, sequence, and genomic location in a comprehensive NoSQL database designed to efficiently support queries and overcome traditional computational and storage limitations associated with analyzing large-scale, highly sparse metagenomic data.

Each MSP carries independent taxonomic annotations from GTDB-Tk 2.4.1 (GTDB release R220) [17] and MetaPhlAn 3 [8], which can disagree due to differences in classification methodology (whole-genome phylogenetic placement versus clade-specific marker genes). To produce a unified taxonomy for cross-system visualizations, we reconciled the two annotations through a consensus approach: both the GTDB-Tk and MetaPhlAn 3 taxon names were mapped to full lineages in the NCBI taxonomy, and the consensus lineage was defined as the set intersection of the two NCBI lineage paths. When no intersection existed, the more specific lineage was retained; when only one annotation was available, it was used directly. Of the 13,795 MSPs, 5,147 (37.3%) carried both annotations; among these, 2,496 (48.5%) agreed at the species level and 2,651 (51.5%) required reconciliation. The remaining MSPs carried only one annotation or none: 8,159 (59.1%) carried only a GTDB-Tk annotation, 37 (0.3%) only a MetaPhlAn 3 annotation, and 452 (3.3%) lacked an NCBI-mappable lineage in either system. A consensus lineage was produced for 11,837 MSPs (85.8%); the remaining 1,958 retained the more specific available annotation or were left unassigned where neither system provided NCBI-mappable taxonomy. This consensus taxonomy is used in all taxonomic visualizations (e.g. Sankey diagrams), while the individual GTDB-Tk and MetaPhlAn 3 annotations remain accessible in the MSP catalog and detail pages.

### 2.2 Implementation and architecture

MetaGEAR Explorer is implemented as a layered web application (Figure 1A). The front end (React 18.2, Material UI 6) communicates with the server via RESTful calls. The server (Python 3.11, FastAPI 0.135.1) is organized into three layers: an API layer handling routing, request validation, and serialization; a service layer coordinating search orchestration, evidence computation, and data extraction pipelines; and a data access layer managing MongoDB queries and per-run workspace directories. An adaptive search job queue dispatches BLASTn and BLASTp (BLAST+ 2.17.0) [18] processes with concurrency control that auto-tunes based on system load, while a background cleanup loop automatically clears workspaces after a 24-hour retention time. By default, hits are filtered at *≥*95% identity, *≥*90% coverage, E-value *≤*0.01, and bit score *≥*0, each adjustable without re-running the underlying alignment. The multi-cohort database is stored in MongoDB 7.0 and comprises collections for gene families (MCGC, *>*33 million entries), protein families (MCPC, *>*23 million entries), metagenomic species pangenome annotations (MSP, 13,795 entries), Pfam domain annotations (17.4 million entries), and 24 per-cohort databases with sample-level abundance and metadata. MetaGEAR Explorer is accessible through both an interactive web interface and documented RESTful API endpoints at https://metagear-explorer.schirmerlab.de, supporting both rapid manual exploration and streamlined programmatic integration into bioinformatics pipelines. The platform follows a privacy-by-design approach: user-submitted sequences are processed transiently and automatically deleted after a set retention period. No personally identifiable information is collected, and only pseudonymous operational analytics are retained. The platform follows GDPR requirements applicable to research-purpose web services hosted in the EU. Comprehensive user documentation is available directly through the platform, including a step-by-step manual, FAQ section, terms of use, and privacy policy.

**Fig. 1.**
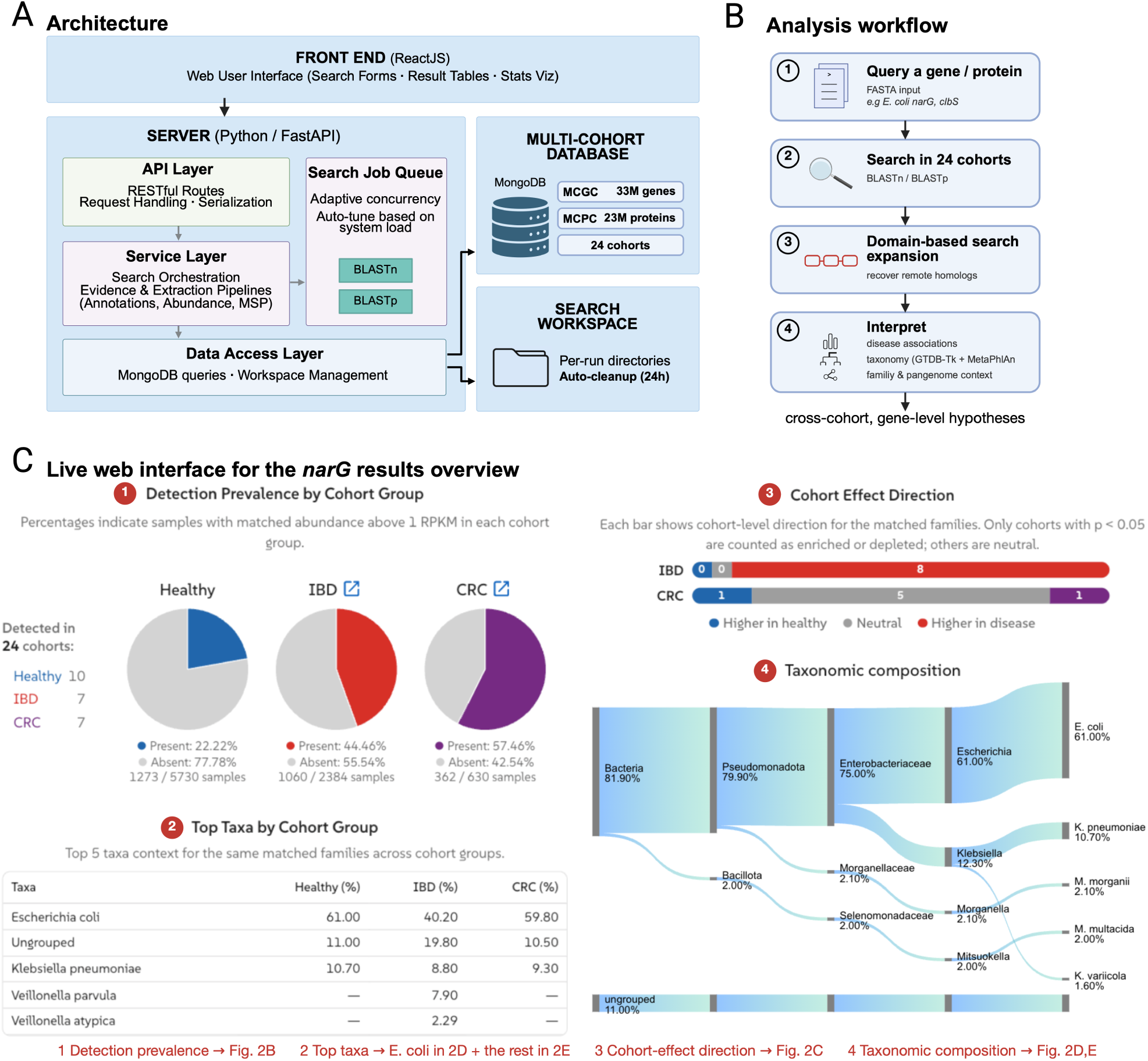
MetaGEAR Explorer platform overview. **(A)** System architecture: a ReactJS front end communicates with a Python/FastAPI server organized into API, service, and data-access layers, backed by a MongoDB multi-cohort database (*>*33M gene families across 24 cohorts); an adaptive job queue dispatches BLAST searches and per-run workspaces are cleared after a 24-hour retention period. **(B)** Analysis workflow: a gene or protein query is searched across the 24 cohorts (BLASTn/BLASTp) and optionally pivoted to its Pfam domain architecture to recover remote homologs, then interpreted through cross-cohort disease association, reconciled taxonomy (GTDB-Tk and MetaPhlAn 3), and family/pangenome context. **(C)** Representative *Results Overview* interface for the *narG* domain search (160 gene families), showing the four linked analytical components: cross-cohort detection prevalence, top taxa, cohort effect direction, and taxonomic composition. Callouts link each interface element to its derived data panel in Figure 2: detection prevalence (2B), top taxa (*E. coli* in 2D, other taxa in 2E), cohort effect direction (2C), and taxonomic composition (2D,E; the full reconciled-taxonomy Sankey is shown in Supplementary Figure 2D); these panels are computed from the same query through the documented public API (accessed June 2026). Full interface views are provided in Supplementary Figures 2 to 5.

### 2.3 Detection prevalence and replication-based cross-cohort enrichment

Regardless of search type, all cross-cohort statistics are computed at the gene-family (MCGC) level: BLASTn matches gene families directly, whereas BLASTp and Pfam-domain searches match protein families in the MCPC that are expanded to their constituent gene families before analysis (a single protein family, clustered at 90% amino-acid identity, can correspond to several gene families clustered at 95% nucleotide identity). Unless stated otherwise, reported family counts therefore refer to gene families. Detection prevalence was defined for each cohort group (Healthy, IBD, and CRC) as the percentage of samples in which a matched gene family reached an abundance above 1 RPKM (reads per kilobase per million mapped reads). Within each cohort, differential abundance was evaluated separately for each gene family by comparing each disease sub-group (Crohn’s disease [CD], ulcerative colitis [UC], or CRC) against that cohort’s control samples with a two-sided

Mann–Whitney U test; a comparison counted as enriched (or depleted) only when it reached *p ≤* 0.05 with a positive (or negative) fold change, and as neutral otherwise. A within-cohort control group is required, so the all-ulcerative-colitis PROTECT cohort contributes to detection but not to the effect-direction consensus, leaving up to six testable IBD and seven CRC cohorts. We did not apply per-feature multiple-testing correction by design: the claim is not that an individual gene is significant within a single study, but that its effect direction reproduces across independent cohorts, for which cross-study replication is itself the multiplicity safeguard. Because per-cohort gene catalogs and MSPs were assembled independently, effect sizes are not directly comparable across cohorts [19]. Cross-cohort evidence was therefore summarized using a vote-counting-by-direction approach [20, 21] in which each cohort contributes equally regardless of sample size: a gene family or MSP received an IBD or CRC label only when it had a testable case–control contrast in at least three cohorts *and* at least 75% of those cohorts showed a significant (*p ≤* 0.05) effect in the same direction. Because the per-cohort catalogs are assembled independently, a false label must arise from the same chance directional signal recurring in three or more separate studies. To estimate how often this could occur by chance, we permuted the disease and control labels within each cohort, re-ran the per-cohort test to measure how often a cohort votes in a given direction by chance, and combined these rates under the same consensus rule (a significant, concordant direction in at least 75% of the supporting cohorts). Requiring at least three supporting cohorts held the false-label rate below 0.1%, versus *≈*18–19% when single-cohort labels were allowed, as most unfiltered labels rest on a single cohort (Supplementary Table 4). These labels are therefore indicators of cross-cohort reproducibility rather than calibrated effect estimates (see Discussion). Statistical analyses were performed in Python 3.11 using SciPy 1.17.1. Unless otherwise stated, all percentages reported in the Results are detection prevalences.

### 2.4 Use of large language models

During the development of MetaGEAR Explorer and the preparation of this manuscript, Claude (Anthropic, Claude Opus 4.8) was used to assist with software engineering tasks and manuscript writing, including spell-checking and text editing. All AI-generated outputs were reviewed, verified, and edited by the authors, who take full responsibility for the content of this work.

## 3 Results

To demonstrate the utility of MetaGEAR Explorer, we present two complementary showcases that exercise the platform’s full feature set. The first is a step-by-step investigation of the *narG* gene that traces a single sequence query through cross-cohort disease association, domain-based expansion, taxonomy reconciliation, and pangenome context (Figure 1B). The second is a programmatic case study of colibactin self-protection (*clbS* / DUF1706) genes that combines web-based discovery with RESTful API multi-variant aggregation, illustrating how the platform scales from interactive hypothesis generation to systematic cross-variant analysis.

### 3.1 Showcase 1: Tracing *narG* from a single sequence to the diversity of nitrate respiration in the inflamed gut

MetaGEAR Explorer provides two complementary gene-centric search modes: sequence-based (BLASTn/BLASTp) and domain-based (using Pfam IDs) searches. Together, they enable researchers to move from searching a specific gene of interest to investigating the full functional diversity of homologs across the multi-cohort database (Figure 1B). Users can paste or upload up to five nucleotide or amino acid sequences at once; the platform auto-detects the input type, searches against the MCGC or MCPC BLAST databases within seconds, and presents alignment statistics (percent identity, sequence coverage, E-value, bit score) with adjustable thresholds on the *Search Results* page (Figure 2A; Supplementary Figure 2A). Alternatively, users can query directly by Pfam domain identifiers to retrieve all protein families sharing a specified domain architecture. The domain pivot capability allows users to transition from the results of a sequence search directly into a domain-based search using the Pfam annotations detected in the matched gene or protein families, bridging sequence similarity and functional domain searches to capture distant homologs that would be missed by strict sequence alignment.

**Fig. 2.**
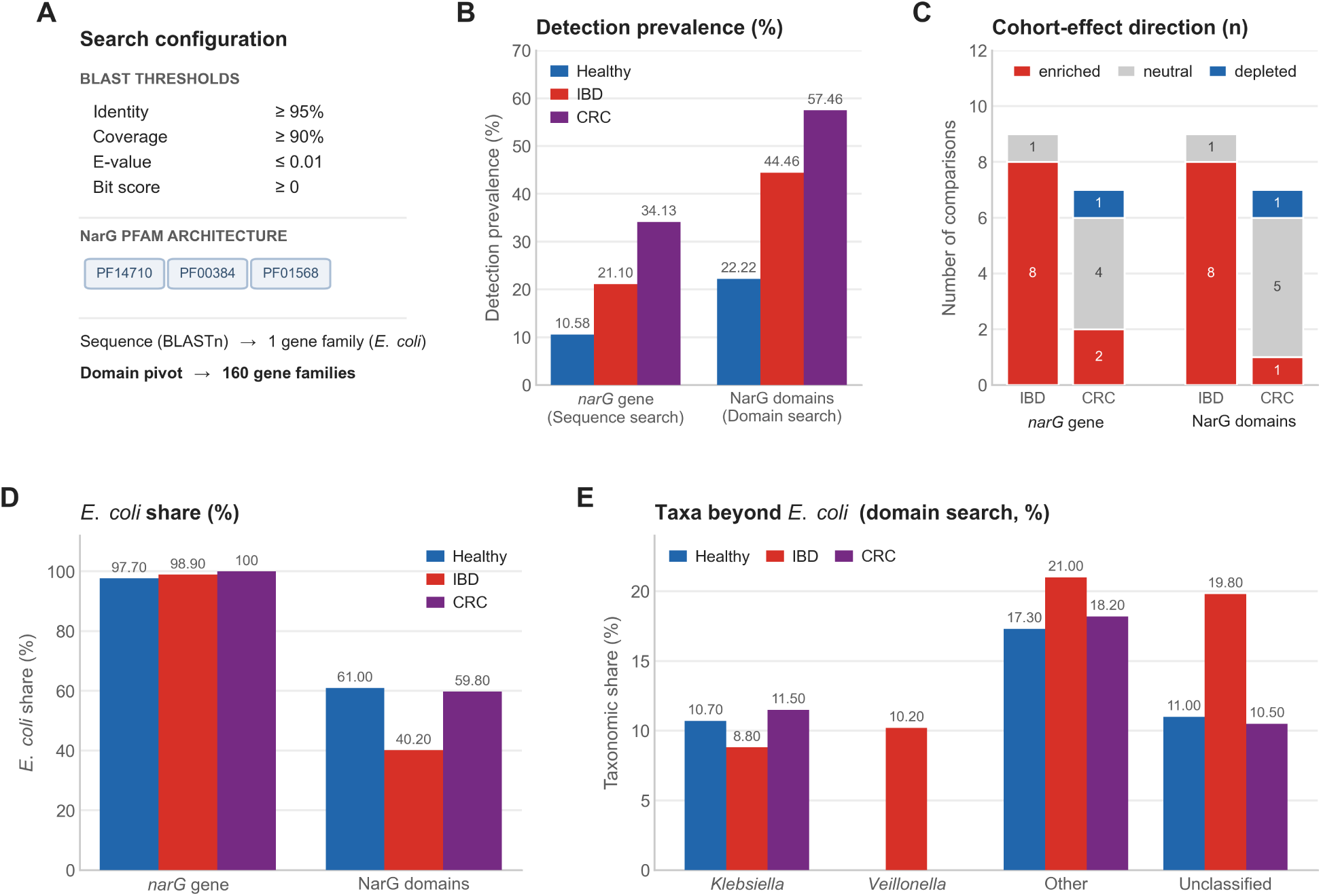
*narG*: sequence search and domain pivot. **(A)** Search configuration: BLAST thresholds and the NarG Pfam domain architecture (PF14710, PF00384, PF01568); the sequence search returns a single *E. coli* family and the domain pivot expands the result set to 160 gene families. **(B)** Detection prevalence by cohort group (Healthy, IBD, CRC) for the sequence search (1 gene family) versus the domain search (160 gene families). **(C)** Cohort-effect direction: for each search and disease group, the number of disease sub-group comparisons that are enriched, neutral, or depleted (two-sided Mann-Whitney U, p *≤* 0.05); a comparison is neutral when it does not reach significance. For IBD, three of the six cohorts contribute separate CD and UC comparisons (nine comparisons in total). **(D)** *E. coli* share of the matched signal: *∼*98–100% for the gene search versus 40–61% for the domain search. **(E)** Domain-search taxa beyond *E. coli* (*Klebsiella*, *Veillonella*, other, and unclassified families) by cohort group; *Veillonella* emerges specifically in IBD. Corresponding interactive interface views are provided in Supplementary Figures 2 and 3.

#### 3.1.1 A single *narG* query expands from one gene family to 160 homologs

To illustrate this workflow, we queried the *E. coli* K-12 *narG* gene (GeneID: 945782), encoding the alpha subunit of respiratory nitrate reductase A [22]. During gut inflammation, host-derived nitric oxide is oxidized to nitrate, providing a respiratory substrate that drives the expansion of *narG* -carrying Enterobacteriaceae [23, 24] and also facilitates ectopic intestinal colonization by oral *Veillonella parvula* via the *nar* operon [25]. While these specific associations are established, we here investigate the full taxonomic repertoire of *narG* - encoding organisms in the diseased gut including distant functional homologs that evade standard sequence-based searches.

A BLASTn search at default thresholds (95% identity, 90% coverage) identified a single gene family with disease-associated prevalence (10.58% Healthy, 21.10% IBD, 34.13% CRC, Figure 2B), driven almost entirely by *E. coli*, whose taxonomic share is 97.70% in Healthy and rises to *∼*98–100% in disease (Figure 2D; Supplementary Figure 1). To capture distant functional homologs missed by this strict alignment, the platform’s domain pivot transitions the query to the NarG Pfam architecture (PF14710, PF00384, and PF01568: the diagnostic domains of membrane-bound respiratory nitrate reductases [22]), expanding the result set from 1 to 160 gene families. This expansion is biologically relevant: beyond *E. coli*, the domain search recovers the genera *Klebsiella* (most prominent in the Healthy and CRC groups) and *Veillonella* (emerging specifically in IBD), shown as genus-level shares by cohort group in Figure 2E. The reconciled-taxonomy Sankey resolves these genera to individual species: *Klebsiella pneumoniae* (10.70% in Healthy), *Veillonella parvula* (7.90% in IBD), and *Veillonella atypica* (2.30% in IBD) (Figure 1C; IBD-filtered view in Supplementary Figure 3C). These organisms, whose nitrate reductase activity in the inflamed gut was recently demonstrated through independent experimental work [25], share insufficient sequence similarity with *E. coli narG* to be detected by a sequence alignment search.

The *Results Overview* page summarizes the matched gene families through several linked analytical components: cross-cohort disease association, functional domain architecture, and taxonomic composition, together with the *Gene/Protein Family Explorer* for inspecting individual gene and protein families. A representative view is shown in Figure 1C, and the full interface in Supplementary Figure 2. Continuing with the NarG domain search (160 gene families), we describe each of these below.

#### 3.1.2 Domain-expanded search uncovers reproducible, cross-cohort IBD enrichment

For each search, MetaGEAR Explorer computes the detection prevalence of matched gene families across cohort groups (Healthy, IBD, and CRC), defined as the percentage of samples with matched abundance above 1 RPKM. For the 160 NarG gene families, detection prevalence rises from 22.22% in the Healthy group to 44.46% in IBD and 57.46% in CRC, consistent with the established role of nitrate respiration in driving Enterobacteriaceae expansion in the inflamed gut [23, 24] (Figure 2B; Supplementary Figure 2B). To assess cross-cohort reproducibility, a cohort effect direction summary quantifies how many independent cohorts show significantly higher abundance in disease versus healthy samples (two-sided Mann-Whitney U test, p *≤* 0.05). For the NarG gene families, this reveals enrichment across the IBD cohorts: of the nine sub-group comparisons spanning the six IBD cohorts with a within-cohort control (three of which split into separate CD and UC sub-groups), all eight that reached significance were enriched and none were depleted (Figure 2C; Supplementary Table 3). This consistent enrichment across diverse cohorts indicates that nitrate reductase expansion is a reproducible feature of IBD dysbiosis rather than a cohort-specific association. By contrast, the CRC signal is more heterogeneous, with cohorts reaching significance in opposing directions (Figure 2C), suggesting a context-dependent association in colorectal cancer. Because per-cohort gene catalogs and MSPs are assembled independently, direct pooling of effect sizes would be inappropriate [19]; instead, our platform employs a vote-counting-by-direction approach [20, 21]: only labels supported by at least three cohorts with *≥*75% concordant significant direction are displayed, ensuring that annotated associations reflect reproducible cross-cohort signals.

#### 3.1.3 Reconciled taxonomy broadens the signal beyond *E. coli* across multiple nitrate-respiring lineages

The domain architecture section presents Pfam functional domains arranged from N- to C-terminal (Figure 2A; Supplementary Figure 2C). For each domain, detailed information is retrieved in real time from InterPro [26] and UniProt [27], including functional descriptions, protein family classifications, and links to AlphaFold [28] structure predictions and Protein Data Bank (PDB) [29] entries. This integration enables researchers to immediately assess whether a matched gene family encodes a complete, well-characterized functional unit or a partial/divergent variant.

Species-level taxonomic composition is visualized as an interactive Sankey diagram using the consensus taxonomy reconciled from GTDB-Tk [17] and MetaPhlAn 3 [8] annotations (see Methods) (Figure 1C; Supplementary Figure 2D), with the cohort-group taxonomic shares summarized in Figure 2D,E. This reconciled taxonomy effectively rescues species-level resolution for MSPs where GTDB-Tk and MetaPhlAn 3 disagree (51.5% of dually annotated MSPs; see Methods), providing a unified taxonomic framework that neither system achieves alone. For the 160 NarG gene families, the Sankey reveals that healthy individuals harbor nitrate reductase homologs across multiple Enterobacteriaceae lineages: *E. coli* (61.00%), *Klebsiella pneumoniae* (10.70%), and ungrouped families (11.00%). This is a substantially broader taxonomic distribution than the single BLAST-matched *E. coli* family (97.70%) and indicates that nitrate respiration is a conserved metabolic strategy across Enterobacteriaceae [23].

#### 3.1.4 Disease stratification: a robust IBD versus a heterogeneous CRC signal

For the NarG domain search, the *Disease Details* pages provide focused analysis of the IBD and CRC associations (Figure 3; Supplementary Figure 3). Gene prevalence is stratified by disease sub-group (e.g., Control, CD, UC for IBD; Figure 3A); a per-cohort breakdown, further resolved by abundance level, is provided in Supplementary Figure 3A. This reveals that the IBD enrichment is consistent across both CD (44.80%) and UC (43.98%) relative to controls (25.09%) within IBD cohorts (Figure 3A). A scatter plot of mean fold change versus significance per cohort comparison (Figure 3B; Supplementary Figure 3B) shows that all IBD cohorts have positive fold-change for *narG*, with the strongest signal in the IBD_Elinav_2022 cohort (fold change 4.07, p = 2.45*×*10*^−^*^11^). In the CRC cohorts, prevalence rises only modestly (57.46% in CRC compared to 52.11% in controls; Figure 3A), and per-cohort comparisons show no consistent direction (Figure 3B), consistent with the weaker, heterogeneous CRC association noted above. Furthermore, disease-filtered taxonomic profiles (cohort-group taxa shares in Figure 2E; the IBD-filtered Sankey in Supplementary Figure 3C) reveal disease-specific taxonomic shifts. For instance, *V. parvula* is unique to IBD, absent from both the Healthy and CRC groups, independently corroborating its demonstrated ability to colonize the inflamed gut by respiring the host-derived nitrate generated during inflammation [25].

**Fig. 3.**
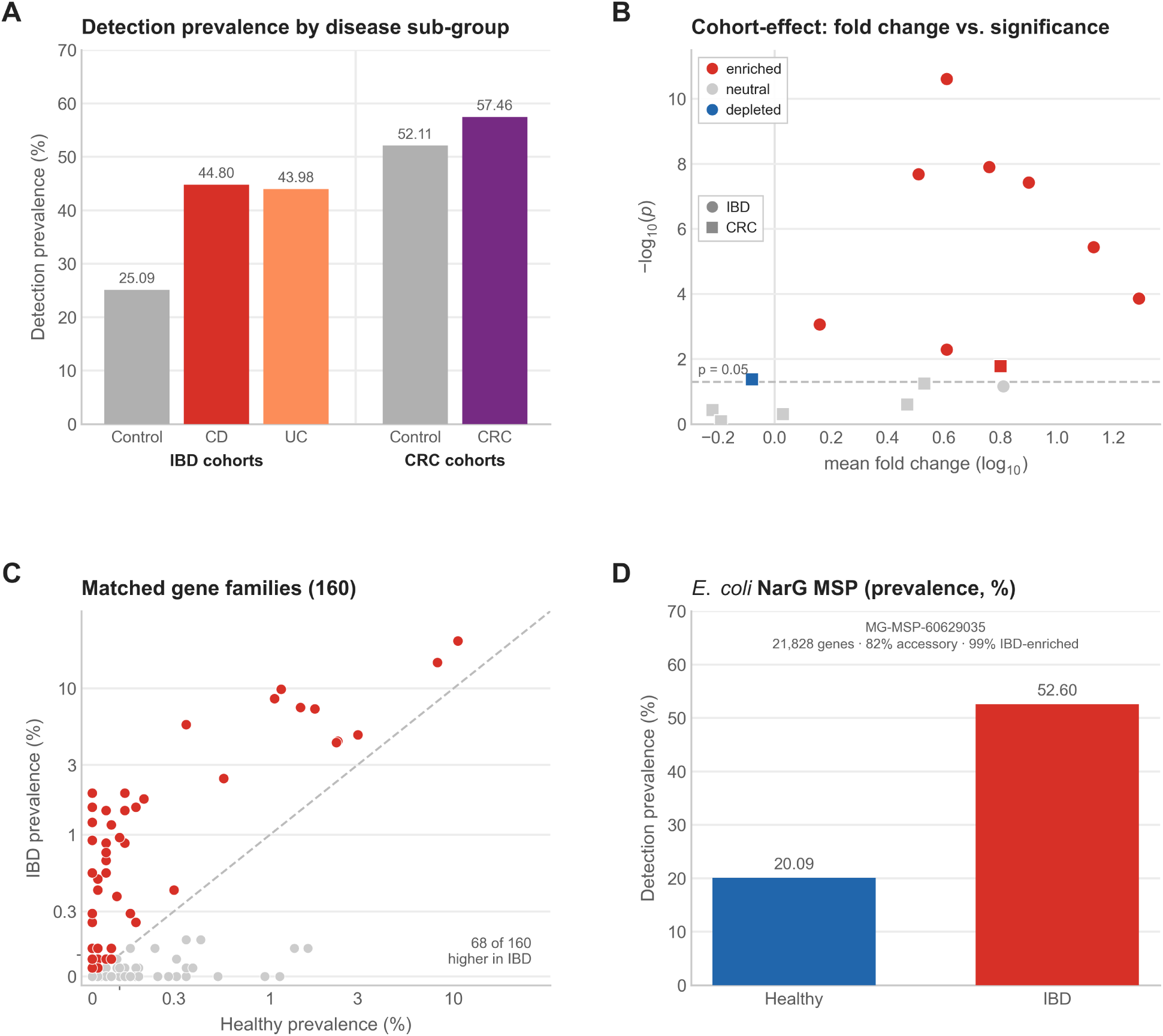
*narG*: disease details and pangenome exploration. All analyses use the NarG domain search (160 gene families). **(A)** Disease sub-group detection prevalence grouped by cohort type (IBD cohorts: Control, CD, UC; CRC cohorts: Control, CRC). **(B)** Cross-cohort association for each cohort/sub-group comparison: mean fold change (log_10_) versus significance (*−*log_10_ p) for IBD (circles) and CRC (squares), coloured enriched, neutral, or depleted (significant at p *≤* 0.05). **(C)** All 160 NarG-domain gene families by Healthy versus IBD prevalence (symlog axes); gene families above the diagonal have higher detection prevalence in IBD than in Healthy. **(D)** Representative *E. coli* NarG MSP (MG-MSP-60629035): disease prevalence and pangenome summary (open pangenome; fraction of genes IBD-enriched). Interactive family- and MSP-level interface views are provided in Supplementary Figures 4 and 5.

#### 3.1.5 Pangenome context places *narG* as a conserved core gene in an intact *nar* operon

The *Gene/Protein Family Explorer* (Supplementary Figure 4A) lets users inspect and prioritize the matched gene and protein families to find those that carry the disease signal. All matched gene and protein families are presented in an interactive, sortable and filterable table, each annotated with its per-group detection prevalence, the number of cohorts in which it was detected, and its MSP (organism) assignment, so users can quickly identify the families that are most prevalent, most reproducible, and taxonomically informative. Those showing reproducible cross-cohort enrichment or depletion additionally carry an “IBD” or “CRC” label from the direction-of-effect consensus described above (a gene or protein family receives this label only when detected in *≥*3 cohorts with *≥*75% concordant significant direction) [30, 31]. Users can then select a subset, preview its combined prevalence and signal coverage, and launch a child overview that recomputes every metric for that subset alone. This is particularly useful when a domain search returns hundreds of gene families but the signal concentrates in a few high-prevalence contributors, which can then be isolated and re-analysed in a single step.

Selecting an individual gene or protein family from the table (Supplementary Figure 4A; the matched gene families are plotted by Healthy-versus-IBD prevalence in Figure 3C) reveals its MSP context (Supplementary Figure 4B), including the consensus taxonomy, gene category within each MSP (core, accessory, or singleton), and cross-cohort disease-association labels. For the *narG* gene family, this reveals its presence across 24 MSPs uniformly classified as *E. coli*, predominantly categorized as a core gene. This indicates that nitrate reductase is a conserved metabolic capability of *E. coli* in the human gut [23, 24], not a strain-variable accessory trait, a distinction critical for determining whether a disease-enriched gene represents a species-wide function or a strain-specific virulence factor.

Users are also able to explore the genomic neighborhood of a gene family, displaying neighboring genes on a representative contig with their predicted functional domains, strand orientation, and relative position; when multiple MSPs are selected, a multi-locus comparison aligns these neighborhoods side by side with anchor-based alignment and ribbon connectors highlighting shared domain architectures (Supplementary Figure 4B). For *narG*, this view is anchored on the NarG domain architecture (PF14710, PF00384, PF01568) and shows a conserved arrangement of flanking genes shared across the *E. coli* MSPs, consistent with an intact *nar* operon context, enabling researchers to identify conserved or divergent operon structures across species without manually aligning contigs.

This gene/protein family view links directly to the interactive MSP Viewer pangenome graph (described in the MSP Catalog Browser section below), bridging search-based and organism-level exploration.

### 3.2 MSP catalog browser: a taxonomy-driven entry point

In addition to the search-based entry points, a standalone MSP catalog browser exposes the full inventory of 13,795 MSPs across all 24 cohorts as a sortable, searchable table listing each MSP with its identifier, cohort of origin, disease group, consensus taxonomy (see Methods), pangenome size, and prevalence (Supplementary Figure 5A). MSPs with a predominant cross-cohort enrichment or depletion are flagged with “IBD” or “CRC” labels, and users can filter by broad taxonomy (phylum, family, and genus), disease group, minimum gene count, and prevalence, or free-text search across identifiers and taxonomic labels for a specific species of interest (e.g., *Fusobacterium nucleatum*). This taxonomy-driven entry point is particularly useful for organism-level hypothesis generation without formulating a prior sequence or domain query.

Selecting any MSP opens a detail page (Supplementary Figure 5B) presenting its consensus taxonomy from GTDB-Tk and MetaPhlAn 3, an associated *R*^2^ score for the linear fit between the MSP abundance profile and the MetaPhlAn 3 species profile across samples [15], a gene composition breakdown into core, accessory, and singleton categories (as defined by MSPminer [16]), and disease prevalence bars for the originating cohort; Figure 3D illustrates a representative *E. coli* NarG MSP (MG-MSP-60629035). The page also embeds an MSP Viewer, an interactive pangenome graph of the gene co-localization network (Supplementary Figure 5C), in which node fill encodes disease association (red, IBD-enriched; purple, CRC-enriched; blue, depleted; gray, neutral) and rings denote gene category (teal, core; amber, accessory; gray, singleton), with edges connecting adjacent genes that co-occur on assembled contigs (weighted by the number of supporting contigs). The disease-association node labels derive from per-gene-family differential abundance tests (two-sided Mann-Whitney U, p *≤* 0.05) across all disease cohorts in which the gene family was detected, consolidated via the MCGC representative mapping and retained only where the family was testable in *≥*3 cohorts and *≥*75% agreed on a significant direction. An interactive dropdown toggles these markers by disease type and direction, revealing genomic regions where disease-associated families cluster; for example, in *E. coli* MSPs, *narG* and neighboring nitrate reductase operon genes appear as co-localized IBD-enriched nodes (Supplementary Figure 5C).

### 3.3 Data export and programmatic (API) access

To support reproducibility and collaboration, MetaGEAR Explorer offers multiple export formats, including CSV evidence summaries and a self-contained JSON export that captures the complete search context and can be re-imported via the Load results tab to restore a full interactive session without server recomputation. Beyond the web interface, all functionality is accessible through documented RESTful API endpoints (OpenAPI specification at https://metagear-explorer.schirmerlab.de/api/docs; Supplementary Table 2); whereas the web interface is optimized for rapid, interactive hypothesis generation, as demonstrated by the *narG*workflow above, the programmatic API enables systematic, multi-query analyses that scale efficiently across dozens or hundreds of queries.

### 3.4 Showcase 2: Colibactin immunity genes reveal a two-layered disease ecology

Having demonstrated guided, web-based exploration with *narG*, we next extend an initial web-based discovery through scripted API calls to aggregate results across multiple ClbS sequence variants, revealing disease-associated patterns that emerge only when the variants are analyzed together.

The genotoxin colibactin, synthesized by a 19-gene biosynthetic cluster on the *pks* genomic island in certain *E. coli* strains, is a microbial virulence factor implicated in CRC tumorigenesis [32]. The *pks* island also encodes self-protection through the *clbS* gene, which neutralizes colibactin autotoxicity, and structurally related ClbS-like proteins have been identified in non-producing bacteria, where they confer immunity against colibactin-triggered prophage induction [33, 34]. Despite growing interest in the ecological role of colibactin immunity, the distribution of ClbS homologs across large-scale gut metagenome collections has not been systematically investigated. MetaGEAR Explorer was used to address this gap across 9,053 gut metagenomes.

#### 3.4.1 Sequence and domain searches reveal disease-enriched *clbS* within a depleted healthy-associated DUF1706 pool

The amino acid sequences of two canonical ClbS protein variants (UniRef100 Q0P7K8 and A0A444RFV0; Supplementary Material) were queried via BLASTp at default thresholds (95% identity, 90% coverage). This identified two matched gene families detected across 20 of 24 cohorts (Supplementary Figure 6A). The absolute prevalence was low, consistent with the specialized nature of colibactin-producing *E. coli*, which constitute a minority of gut Enterobacteriaceae even in disease. However, a clear disease-associated expansion was observed: 1.85% prevalence in healthy controls versus 6.21% in IBD and 7.62% in CRC (Figure 4A). Even at low prevalence, this enrichment is biologically relevant: colibactin-producing strains exert disproportionate genotoxic effects through DNA alkylation and double-strand breaks [32], and their expansion is particularly notable in UC, where chronic colonic inflammation is a known driver of colorectal carcinogenesis and may link colibactin-producing *E. coli* to the elevated CRC risk observed in UC patients [5]. Taxonomic profiling showed that the signal was driven almost entirely by *E. coli* (94.40% in Healthy and IBD, 69.90% in CRC; Figure 4C). The cohort effect direction summary indicated that where significant abundance changes occurred within individual IBD cohorts (2 of 6 cohorts in which ClbS was detected), they consistently showed disease enrichment without any contradictory depletion (two-sided Mann-Whitney U test, p *≤* 0.05; Supplementary Figure 6A). Within IBD, the enrichment was strongest in UC (8.06%) compared to CD (4.91%), aligning with the reported expansion of Enterobacteriaceae and other Proteobacteria in ulcerative colitis [24, 35].

**Fig. 4.**
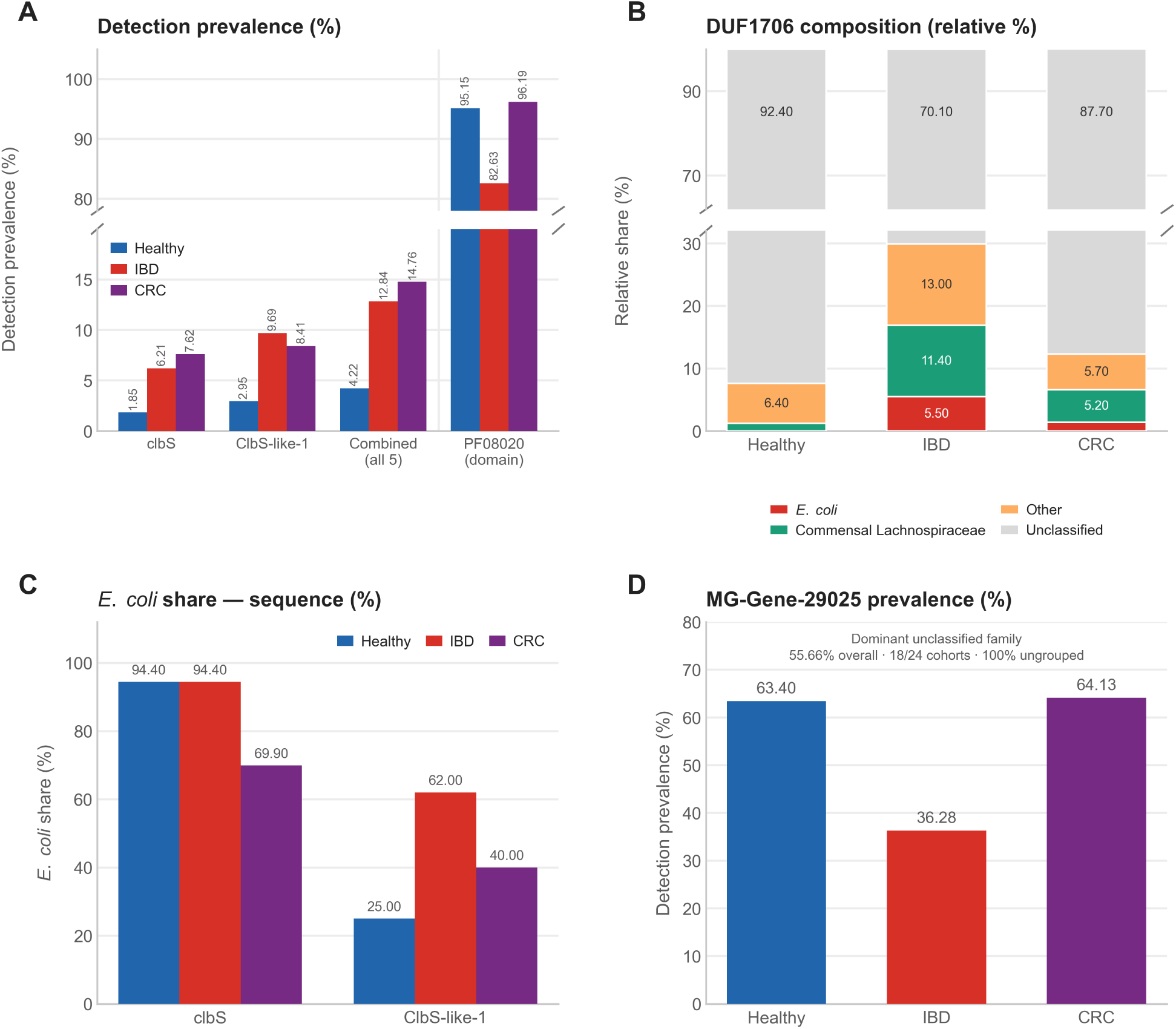
Colibactin immunity genes reveal a two-layered disease ecology. **(A)** Detection prevalence by cohort group for canonical *clbS*, ClbS-like-1, the combined five-variant set, and the PF08020 (DUF1706) domain family (note the broken y-axis): sequence-based searches rise in disease while the broad domain family is depleted in IBD. **(B)** DUF1706 taxonomic composition by cohort group (relative shares): the unclassified, healthy-associated majority recedes in IBD as the *E. coli* share rises. **(C)** *E. coli* share of the sequence searches: canonical *clbS* is stably at *∼*94% *E. coli*, whereas ClbS-like-1 shifts from 25% (Healthy) to 62% (IBD). **(D)** The dominant unclassified family MG-Gene-29025 (55.66% overall prevalence, 100% ungrouped taxonomy) drops from 63.40% in Healthy to 36.28% in IBD. The corresponding interactive web-discovery view is shown in Supplementary Figure 6.

To test whether this disease association is specific to *clbS* or extends to the broader DUF1706 domain family, a domain-based search was performed using PF08020 (DUF1706), the single Pfam domain identified in the ClbS hits. This revealed a contrasting ecological pattern (Figure 4A,B; Supplementary Figure 6B): 1,121 gene families were matched across all 24 cohorts, with near-ubiquitous prevalence in healthy controls (95.15%, 5,452 samples) and CRC (96.19%, 606 samples). Unlike the specific ClbS variants, the overall PF08020 domain prevalence was reduced in IBD (82.63%, 1,970 samples), indicating that while *E. coli clbS* expands in disease, the healthy-associated bacteria that carry the majority of DUF1706 homologs are depleted. This divergence reflects a pronounced disease-associated phylogenetic shift: whereas the ClbS signal was taxonomically restricted to *E. coli* (Class Gammaproteobacteria, Phylum Pseudomonadota [Proteobacteria]), the broader PF08020 domain in healthy individuals was carried predominantly by an unclassified gene family together with commensal Lachnospiraceae (Class Clostridia, Phylum Bacillota [Firmicutes]). Among classified taxa, *Lachnospira eligens* was the leading commensal contributor in healthy controls (1.20%; 1.20%–2.10% across the three cohort groups), whereas *Roseburia* species contributed mainly in disease, as the dominant unclassified family recedes: *Roseburia intestinalis* (7.90% in IBD, 1.30% in CRC) and *Roseburia inulinivorans* (1.90% in CRC). *E. coli* appeared only as a minor contributor in disease (5.50% in IBD, negligible in Healthy). This pattern connects the established IBD-associated depletion of butyrate-producing Clostridia and expansion of Gammaproteobacteria [23, 36] with a concurrent loss of healthy-associated DUF1706 homologs whose functional role remains uncharacterized. This dysbiotic shift involving a single domain family suggests that the DUF1706 domain may serve as a molecular marker for tracking the Clostridia-to-Gammaproteobacteria transition at the gene level.

#### 3.4.2 Programmatic variant aggregation uncovers hidden prevalence and a *Proteus mirabilis* carrier

To further characterize the ClbS disease enrichment observed in the web-based analysis, MetaGEAR Explorer’s RESTful API was used to systematically query five ClbS and ClbS-like protein sequences (identified by Mäklin et al. [33]): the canonical ClbS (Q0P7K8), a ClbS variant (A0A444RFV0), and three ClbS-like proteins (A0A376RIA6, A0A2C5TQB2, A0A631IX87). The canonical ClbS variants (Q0P7K8 and A0A444RFV0) mapped to the same gene family (MG-Gene-43344), confirming that the single-residue difference between them falls within the clustering threshold. The ClbS-like-1 protein (A0A376RIA6) exhibited the highest disease prevalence of all queries, enriched in both IBD (9.69%) and CRC (8.41%) compared to healthy controls (2.95%). Disease-stratified taxonomic profiling revealed a compositional shift: in healthy individuals, only 25.00% of the ClbS-like-1 signal was attributable to *E. coli*, but in IBD this fraction rose to 62.00%, a 2.5-fold enrichment of the *E. coli* contribution specifically in disease samples (Figure 4C).

MSP analysis additionally identified *Proteus mirabilis* as a carrier of ClbS-like homologs in two independent pangenomes, detected predominantly in IBD samples (0.92% IBD vs. 0.07% Healthy). Although the absolute detection frequency is low, this observation is notable because *P. mirabilis* has been independently established as a pathobiont in both Crohn’s disease [37] and ulcerative colitis [35], capable of inducing colonic inflammation in animal models. Determining whether the *P. mirabilis* DUF1706 homolog retains the colibactin-neutralizing activity demonstrated for other non-producer ClbS variants [34] represents an experimentally testable hypothesis generated directly from this metagenomic analysis.

The remaining ClbS-like variants (A0A2C5TQB2, A0A631IX87) produced no matches at the stringent default thresholds, indicating that these more divergent DUF1706 homologs lack close paralogs in the gut metagenome catalog. Aggregating all five queries yielded a combined prevalence of 4.22% in healthy controls versus 12.84% in IBD and 14.76% in CRC (Figure 4A). This programmatic approach enabled per-variant comparison and aggregation that would be impractical through the web interface alone, demonstrating that tracking isolated sequence variants substantially underestimates the aggregate prevalence of colibactin immunity genes.

#### 3.4.3 The two layers mirror an IBD shift from butyrate-producing commensals to colibactin-immune Gammaproteobacteria

Together, the sequence and domain searches reveal a two-layered ecological dynamic (Figure 4). The DUF1706 domain (PF08020) is widespread, carried by *>*95% of healthy gut microbiomes, predominantly by an unclassified gene family together with commensal Lachnospiraceae such as *Lachnospira eligens*. Butyrate-producing *Roseburia* species, whose depletion characterizes IBD dysbiosis [36], form a larger share of this domain pool in disease as the unclassified majority recedes. Within this broader family, a functionally distinct subset has been characterized as colibactin self-protection determinants: ClbS neutralizes colibactin autotoxicity in *pks*-carrying *E. coli*, while ClbS homologs in non-producing bacteria provide immunity against colibactin-triggered prophage induction [34].

In IBD, this equilibrium shifts in two directions. Commensal DUF1706 carriers are depleted, with PF08020 prevalence dropping from 95.15% to 82.63%. Simultaneously, *E. coli* lineages carrying the colibactin-specific *clbS* expand from 1.85% to 6.21%. This compositional change is directly evidenced by the disease-conditioned taxonomic profile of ClbS-like-1, where the *E. coli* contribution rises from 25.00% in healthy individuals to 62.00% in IBD (Figure 4C).

Beyond *E. coli*, ClbS-like homologs were independently detected in *Proteus mirabilis*, carried in two separate pangenomes almost exclusively in IBD samples, consistent with recent findings that colibactin immunity genes are distributed across diverse Gammaproteobacteria [33, 34]. This observation fits the broader dysbiotic pattern in IBD: the depletion of commensal Bacillota reduces butyrate production and anti-inflammatory signaling [36], potentially favoring the expansion of Gammaproteobacteria equipped with colibactin immunity. Notably, a single uncharacterized gene family (MG-Gene-29025) accounts for over 60% of the total PF08020 signal and is present in 63.40% of healthy samples but only 36.28% in IBD (Figure 4D; Supplementary Figure 7A,B). This family has no MSP assignment and its taxonomy remains entirely ungrouped, suggesting it represents a widespread healthy-associated element depleted during IBD dysbiosis.

## 4 Discussion

MetaGEAR Explorer fills a gap in the current landscape of metagenomic analysis tools (Table 1). While functional profiling tools such as HUMAnN 3 quantify gene family and pathway abundances against curated reference databases, and de novo catalogs such as IGC and UHGP provide downloadable gene inventories without disease context, MetaGEAR Explorer combines a de novo assembled gene catalog with built-in cross-cohort disease stratification and interactive exploration, all accessible through both a web interface and a programmatic API.

A key strength lies in the cross-cohort evidence framework. Rather than reporting associations from a single study, the platform quantifies the direction and consistency of disease associations across all available cohorts, helping researchers distinguish reproducible signals from cohort-specific associations. Because per-cohort gene catalogs and MSPs are assembled independently, each capturing the specific sequence diversity of its cohort, effect sizes are not directly poolable across studies. The platform therefore employs a vote-counting-by-direction approach [20] that leverages this independence as a strength: concordant signals across independently assembled cohorts provide inherently robust evidence of reproducibility, as each cohort serves as an independent validation [19]. For *narG*, this framework reveals consistent IBD enrichment across cohorts (eight of the nine sub-group comparisons across six cohorts significantly enriched, none depleted), a cross-cohort consistency that complements the mechanistic evidence from individual studies [23] by demonstrating the reproducibility of this association across diverse and geographically distinct patient populations. The CRC signal, by contrast, is heterogeneous, reflecting the known biological variability across CRC studies [7, 38]. This distinction between robust and heterogeneous signals is precisely what multi-cohort evidence is designed to reveal. Together, these results extend the taxonomic repertoire of nitrate-respiring organisms in the inflamed gut beyond the canonical Enterobacteriaceae-centered model, and demonstrate that domain-aware searches can recover functionally relevant homologs (e.g., *Veillonella* spp.) that strict sequence alignment misses.

The cross-cohort evidence is complemented by gene-family-level exploration features (MSP assignments and genomic neighborhood context) that enable researchers to move from a statistical association to a mechanistic hypothesis. For example, the *narG* MSP context shows that nitrate reductase is a conserved core gene across *E. coli* pangenomes, while the genomic neighborhood view is consistent with an intact *nar* operon context, together indicating that the disease-enriched signal reflects a conserved metabolic capability rather than a strain-variable accessory trait. The ability to transition from sequence to domain-based searches further extends this by testing whether enrichment is driven by a single gene family or by the broader biological function encoded by its domain architecture, as demonstrated by the expansion from 1 *narG* BLAST hit to 160 NarG-domain gene families including several *Veillonella* species. The MSP catalog browser provides a complementary, taxonomy-driven entry point for organism-level hypothesis generation, and the MSP Viewer brings interactive pangenome visualization to a web environment that requires no local compute infrastructure or specialized bioinformatics expertise [15].

The documented RESTful API enables systematic, multi-query analyses beyond what manual exploration supports. The colibactin case study illustrates this: an initial web-based discovery was extended through scripted API calls to aggregate five ClbS/ClbS-like variants, revealing that their combined prevalence (12.84% IBD, 14.76% CRC) substantially exceeds any single variant alone. Self-contained JSON exports that can be re-imported for offline viewing further support transparency and reproducibility.

Several limitations should be noted. First, the current scope is restricted to the human gut microbiome in the context of IBD and CRC, so associations identified here may not generalize to other body sites, host organisms, or disease phenotypes. Second, the underlying multi-cohort database is pre-computed and static; users querying recently published cohorts not yet integrated into the catalog will not find them represented in cross-cohort summaries until the next release. Third, the disease enrichment labels rest on cross-cohort directional concordance (a significant effect in the same direction in at least 75% of the *≥*3 testable cohorts) rather than per-feature multiple-testing correction; replication controls chance (empirical false-discovery rate *≈*0.05–0.07%) but not correlated technical bias, since cohorts that share sequencing platforms or reference databases can reproduce the same artefact [19]. Fourth, the contrast between the robust IBD signal and the more heterogeneous CRC signal partly reflects statistical power: the CRC groups are considerably smaller (630 versus 2,384 samples), so a weaker or absent CRC consensus does not necessarily indicate weaker underlying biology.

## 5 Conclusions

Cross-cohort functional meta-analysis with MetaGEAR Explorer recovers reproducible, biologically interpretable microbial gene signatures of gastrointestinal disease across 9,053 metagenomes. The *narG* walkthrough illustrates how a sequence-to-domain pivot expands a single BLAST hit into 160 functional homologs with consistent, reproducible IBD enrichment across cohorts, including *Veillonella parvula* and *V. atypica* that strict sequence alignment alone misses. The colibactin case study demonstrates how programmatic aggregation across multiple sequence variants exposes an opposing two-layered ecology, in which canonical *clbS* expansion in disease coexists with depletion of the broader DUF1706 domain family carried by commensal Clostridia and a dominant unclassified, healthy-associated family, and surfaces *Proteus mirabilis* as a ClbS-like carrier in IBD not previously highlighted in large-scale gut-metagenome cohorts.

Together, these worked examples establish MetaGEAR Explorer as a platform that bridges catalog-scale metagenomic data and the generation of testable functional hypotheses of the role of the microbiome in disease, enabling effective cross-cohort functional meta-analysis within seconds without local computational resources or specialized bioinformatics expertise. Future releases will broaden the platform to additional body sites, host organisms, and disease phenotypes, integrate newly published cohorts as the underlying database is periodically re-processed, and support user-uploaded datasets for direct comparison against the existing catalog.

## List of abbreviations

API: Application Programming Interface
BLAST: Basic Local Alignment Search Tool
CD: Crohn’s disease
cMD3: curatedMetagenomicData version 3
CRC: Colorectal cancer
CSV: Comma-Separated Values
DUF: Domain of Unknown Function
GTDB-Tk: Genome Taxonomy Database Toolkit
HMM: Hidden Markov Model
IBD: Inflammatory bowel disease
IGC: Integrated Gene Catalog
JSON: JavaScript Object Notation
MCGC: Multi-Cohort Gene Catalog
MCPC: Multi-Cohort Protein Catalog
MSP: Metagenomic Species Pangenome
PDB: Protein Data Bank
Pfam: Protein Families database
REST: Representational State Transfer
RPKM: Reads Per Kilobase per Million mapped reads
UC: Ulcerative colitis
UHGG: Unified Human Gastrointestinal Genome
UHGP: Unified Human Gastrointestinal Protein

## Supplementary information

Supplementary tables, figures, example use cases, and test input data are provided in the Appendices.

## Declarations

### Ethics approval and consent to participate

Not applicable.

### Consent for publication

Not applicable.

### Availability of data and materials

The MetaGEAR Explorer web platform is freely available at https://metagear-explorer.schirmerlab.de. Content is distributed under the Technical University of Munich Terms of Use (https://www.mls.ls.tum.de/en/mdi/impressum) and the MetaGEAR Explorer Terms of Use (https://metagear-explorer.schirmerlab.de/terms). All metagenomic sequencing data analyzed in this study were obtained from the 24 previously published cohorts listed in Supplementary Table 1; the original repository accessions (SRA, ENA, EGA, or equivalent) are reported in the corresponding cohort publications cited in this table. The multi-cohort gene catalog (MCGC, *>*33 million non-redundant gene families), multi-cohort protein catalog (MCPC, *>*23 million non-redundant protein families), and metagenomic species pangenome (MSP) collection (13,795 MSPs) underlying MetaGEAR Explorer are described and made available through the accompanying study [15]. Pfam domain annotations are available from InterPro [26]; representative protein structures and functional annotations are retrieved from AlphaFold [28], the Protein Data Bank [29], and UniProt [27]. All quantitative data figures in this study (Figures 2–4) were generated by open-source scripts that query the MetaGEAR Explorer public REST API and render the retrieved data. These scripts, the raw API responses used as figure inputs, the figure-generation code, and a table mapping each figure to the script and data that reproduce it are openly available at https://github.com/schirmer-lab/metagear-explorer-ext and archived at Zenodo (DOI: 10.5281/zenodo.21385398). Analysis code is released under the MIT License and the derived data files under the Creative Commons CC-BY-4.0 License.

The MetaGEAR Explorer platform is described as follows:

- Project name: MetaGEAR Explorer
- Project home page: https://metagear-explorer.schirmerlab.de
- API documentation (OpenAPI): https://metagear-explorer.schirmerlab.de/api/docs
- Operating system(s): Platform-independent (accessed through any modern web browser)
- Programming language: Python (back end); JavaScript / TypeScript (front end)
- Other requirements: FastAPI, MongoDB, BLAST+, React 18, Material UI
- License (analysis code): MIT
- Any restrictions to use by non-academics: None; the web service and REST API are freely accessible without registration.

MetaGEAR Explorer is provided as a maintained, freely accessible hosted service with a documented public REST API, and all analyses reported here are fully reproducible against this API using the archived scripts above. The production deployment source code is specific to the institutional hosting environment and to the GDPR-compliant operation of a public sequence-search service, and is therefore not publicly released; it will be provided to the handling editor and peer reviewers for evaluation during the review process, and can be shared with academic researchers for non-commercial use under an appropriate data-protection and hosting agreement.

### Authors’ contributions

E.R. and S.J. contributed to study design, coordination, software development and testing; C.Z., X.H., F.N., and S.W. contributed to software development and testing. M.S. is responsible for conceptualization, study design, coordination and supervision of all aspects of this project and acquired funding. E.R., S.J., and M.S. wrote the manuscript. All authors reviewed the manuscript.

## Acknowledgements

The authors gratefully acknowledge the Leibniz Supercomputing Centre for providing computing time and support on its Linux-Cluster.

For the LifeLines-DEEP cohort, which was included in this study, we thank the participants and the staff for their contribution; funding for this cohort was provided by the Top Institute Food and Nutrition Wageningen grant GH001, and sequencing was carried out in collaboration with the Broad Institute.

Figures were created with BioRender.com (Rios, E., 2026).

The authors would also like to thank Xin Yin for the cover image design of the MetaGEAR Explorer website.

## Authors’ information

Not applicable.

## Appendix A Supplementary Tables

**Supplementary Table 1:** Metagenomic cohorts included in the Multi-cohort Database used by MetaGEAR Explorer.

| Project_name | Study_name | Ref. |
| --- | --- | --- |
| CRC_GuptaA_2019 | Association of *Flavonifractor plautii*, a Flavonoid-Degrading Bacterium, with the Gut Microbiome of Colorectal Cancer Patients in India | [39] |
| CRC_HanniganGD_2017 | Diagnostic Potential and Interactive Dynamics of the Colorectal Cancer Virome | [40] |
| CRC_LiuNN_2020 | Multi-kingdom microbiota analyses identify bacterial–fungal interactions and biomarkers of colorectal cancer across cohorts | [41] |
| CRC_ThomasAm_2022 | Metagenomic analysis of colorectal cancer datasets identifies cross-cohort microbial diagnostic signatures and a link with choline degradation | [42] |
| CRC_YachidaS_2019 | Metagenomic and metabolomic analyses reveal distinct stage-specific phenotypes of the gut microbiota in colorectal cancer | [7] |
| CRC_YangY_2021 | Dysbiosis of human gut microbiome in young-onset colorectal cancer | [38] |
| CRC_YuJ_2015 | Metagenomic analysis of faecal microbiome as a tool towards targeted non-invasive biomarkers for colorectal cancer | [43] |
| Healthy_500FG | Linking the Human Gut Microbiome to Inflammatory Cytokine Production Capacity | [44] |
| Healthy_DeFilippisF_2019 | Distinct Genetic and Functional Traits of Human Intestinal *Prevotella copri* Strains Are Associated with Different Habitual Diets | [45] |
| Healthy_DhakanDB_2019 | The unique composition of Indian gut microbiome, gene catalogue, and associated fecal metabolome deciphered using multi-omics approaches | [46] |
| Healthy_HansenLBS_2018 | A low-gluten diet induces changes in the intestinal microbiome of healthy Danish adults | [47] |
| Healthy_KeohaneDM_2020 | Microbiome and health implications for ethnic minorities after enforced lifestyle changes | [48] |
| Healthy_LifeLinesDEEP | Cohort profile: LifeLines DEEP, a prospective, general population cohort study in the northern Netherlands: study design and baseline characteristics | [49] |
| Healthy_LokmerA_2019 | Use of shotgun metagenomics for the identification of protozoa in the gut microbiota of healthy individuals from worldwide populations with various industrialization levels | [50] |
| Healthy_MehtaRS_2018 | Stability of the human faecal microbiome in a cohort of adult men | [51] |
| Healthy_PasolliE_2019 | Extensive Unexplored Human Microbiome Diversity Revealed by Over 150,000 Genomes from Metagenomes Spanning Age, Geography, and Lifestyle | [52] |
| Healthy_PREDICT1_2021 | Microbiome connections with host metabolism and habitual diet from 1,098 deeply phenotyped individuals | [53] |
| IBD_Elinav_2022 | Targeted suppression of human IBD-associated gut microbiota commensals by phage consortia for treatment of intestinal inflammation | [54] |
| IBD_HallAB_2017 | A novel *Ruminococcus gnavus* clade enriched in inflammatory bowel disease patients | [55] |
| IBD_HeQ_2017 | Two distinct metacommunities characterize the gut microbiota in Crohn’s disease patients | [56] |
| IBD_HMP2 | Multi-omics of the gut microbial ecosystem in inflammatory bowel diseases | [2] |
| IBD_IjazUZ_2017 | The distinct features of microbial 'dysbiosis' of Crohn's disease do not occur to the same extent in their unaffected, genetically-linked kindred | [57] |
| IBD_LewisJD_2015 | Inflammation, Antibiotics, and Diet as Environmental Stressors of the Gut Microbiome in Pediatric Crohn's Disease | [58] |
| IBD_PROTECT_2022 | Linking microbial genes to plasma and stool metabolites uncovers host-microbial interactions underlying ulcerative colitis disease course | [59] |

**Supplementary Table 2:** MetaGEAR Explorer RESTful API: overview of public endpoint groups. Endpoints are relative to the base URL https://metagear-explorer.schirmerlab.de/api. This table summarizes the main capability groups; the complete, always-current specification (parameters, schemas, and example requests) is available through the interactive Swagger UI at https://metagear-explorer.schirmerlab.de/api/docs.

| Group | Representative endpoints | Description |
| --- | --- | --- |
| Search | POST /search, GET /search/{id}, PUT .../filters | Submit a BLASTn, BLASTp, or Pfam-domain search; poll its status; and adjust alignment thresholds (identity, coverage, E-value, bit score) without re-running BLAST. |
| Results | .../matches, POST /extraction, .../evidence-summary, .../details/{disease} | Retrieve matched gene and protein families with their per-group prevalence, taxonomy, Pfam domains, disease sub-group details, and the cross-cohort evidence summary, including for user-selected subsets. |
| Family | .../families/{...}/msp-context, .../neighborhood | Per-family MSP assignment and consensus taxonomy, and the genomic-neighborhood context of a selected gene family. |
| MSP catalog | /explore/msp/catalog, /explore/msp/{accession}/detail, .../pangenome, .../sequences.fasta | Browse and filter the full MSP catalog (by taxonomy, cohort, gene count, and prevalence), and retrieve MSP detail, the pangenome graph, and gene sequences (FASTA). |
| Export | .../export.json, .../evidence-summary.csv, /media | Download a self-contained, re-importable JSON session, CSV evidence summaries, result-plot images, and a ZIP results bundle. |

**Supplementary Table 3:** Per-cohort case–control comparisons underlying the NarG domain-search cohort-effect summary (Figure 2C) and the fold-change scatter (Figure 3B). Each row is one disease sub-group tested against that cohort’s within-cohort control samples with a two-sided Mann–Whitney U test on gene-family abundance; a comparison is labelled enriched (or depleted) when *p ≤* 0.05 with a positive (or negative) fold change, and neutral otherwise. Mean fold changes are shown on a log_10_ scale as in Figure 3B (positive = higher abundance in disease; for example, the IBD_Elinav_2022 CD value of +0.61 corresponds to a 4.07-fold enrichment). The all-ulcerative-colitis IBD_PROTECT_2022 cohort has no within-cohort control and is excluded from the effect-direction analysis (it contributes to detection prevalence only). Across the nine IBD comparisons (six cohorts, three of which contribute separate CD and UC sub-groups), eight are significantly enriched and none depleted; the seven CRC comparisons comprise one enriched, one depleted, and five neutral.

| Cohort | Comparison | Mean fold change ($\log_{10}$) | $p$-value | Direction |
| --- | --- | --- | --- | --- |
| *IBD cohorts (disease sub-group vs. within-cohort control)* |  |  |  |  |
| IBD_Elinav_2022 | CD vs. control | +0.61 | $2.45 \times 10^{-11}$ | Enriched |
| IBD_Elinav_2022 | UC vs. control | +0.76 | $1.3 \times 10^{-8}$ | Enriched |
| IBD_HallAB_2017 | CD vs. control | +0.61 | $5.1 \times 10^{-3}$ | Enriched |
| IBD_HallAB_2017 | UC vs. control | +1.13 | $3.6 \times 10^{-6}$ | Enriched |
| IBD_HeQ_2017 | CD vs. control | +0.90 | $3.7 \times 10^{-8}$ | Enriched |
| IBD_HMP2 | CD vs. control | +0.51 | $2.1 \times 10^{-8}$ | Enriched |
| IBD_HMP2 | UC vs. control | +0.16 | $8.7 \times 10^{-4}$ | Enriched |
| IBD_IjazUZ_2017 | CD vs. control | +0.81 | $6.9 \times 10^{-2}$ | Neutral |
| IBD_LewisJD_2015 | CD vs. control | +1.29 | $1.4 \times 10^{-4}$ | Enriched |
| *CRC cohorts (CRC vs. within-cohort control)* |  |  |  |  |
| CRC_GuptaA_2019 | CRC vs. control | +0.80 | $1.7 \times 10^{-2}$ | Enriched |
| CRC_HanniganGD_2017 | CRC vs. control | +0.47 | $2.5 \times 10^{-1}$ | Neutral |
| CRC_LiuNN_2020 | CRC vs. control | -0.19 | $8.3 \times 10^{-1}$ | Neutral |
| CRC_ThomasAm_2022 | CRC vs. control | +0.53 | $5.6 \times 10^{-2}$ | Neutral |
| CRC_YachidaS_2019 | CRC vs. control | +0.03 | $5.0 \times 10^{-1}$ | Neutral |
| CRC_YangY_2021 | CRC vs. control | -0.08 | $4.3 \times 10^{-2}$ | Depleted |
| CRC_YuJ_2015 | CRC vs. control | -0.22 | $3.7 \times 10^{-1}$ | Neutral |

**Supplementary Table 4:** Estimate of how often the cross-cohort enrichment labels could arise by chance. Within each cohort, the disease and control labels were repeatedly shuffled (120 permutations) and the per-cohort test re-run to measure how often a cohort votes for enrichment or depletion by chance; these rates were then combined under the platform’s consensus rule (a significant, concordant direction in at least 75% of the supporting cohorts) to give the expected number of false labels and the corresponding false-discovery rate (FDR) at each minimum number of supporting cohorts. Because most labels are supported by a single cohort, requiring at least three supporting cohorts removes the non-reproducible calls and holds the FDR below 0.1%.

| Disease | Minimum supporting cohorts | Displayed labels | Expected false labels | FDR (%) |
| --- | --- | --- | --- | --- |
| IBD | $\geq 1$ (no floor) | 1,713,375 | 318,365 | 18.58 |
| IBD | $\geq 2$ | 263,574 | 1,259 | 0.48 |
| IBD | $\geq 3$ | 147,284 | 67 | 0.046 |
| CRC | $\geq 1$ (no floor) | 354,178 | 66,348 | 18.73 |
| CRC | $\geq 2$ | 45,547 | 348 | 0.76 |
| CRC | $\geq 3$ | 25,944 | 17 | 0.067 |

## Appendix B Supplementary Figures

**Supplementary Figure 1.**
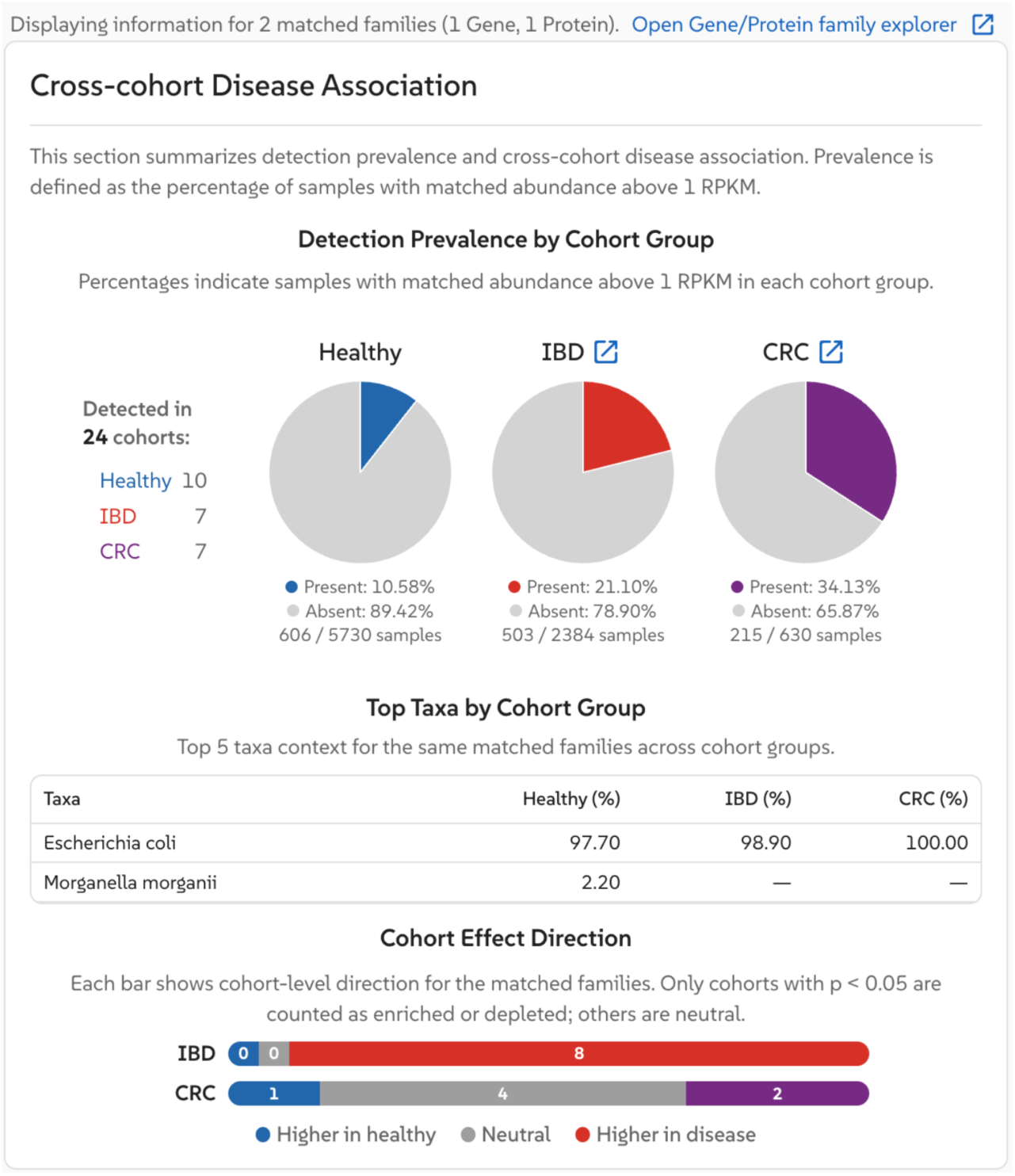
*narG* BLASTn sequence search results. The Results Overview shows detection prevalence by cohort group (10.58% Healthy, 21.10% IBD, 34.13% CRC; 606/5,730, 503/2,384, and 215/630 detected/total samples respectively), top taxa showing *E. coli* as the dominant contributor (97.70% in Healthy), and cohort effect direction summary indicating consistent enrichment in IBD (8 IBD sub-group comparisons significantly higher in disease, none lower).

**Supplementary Figure 2.**
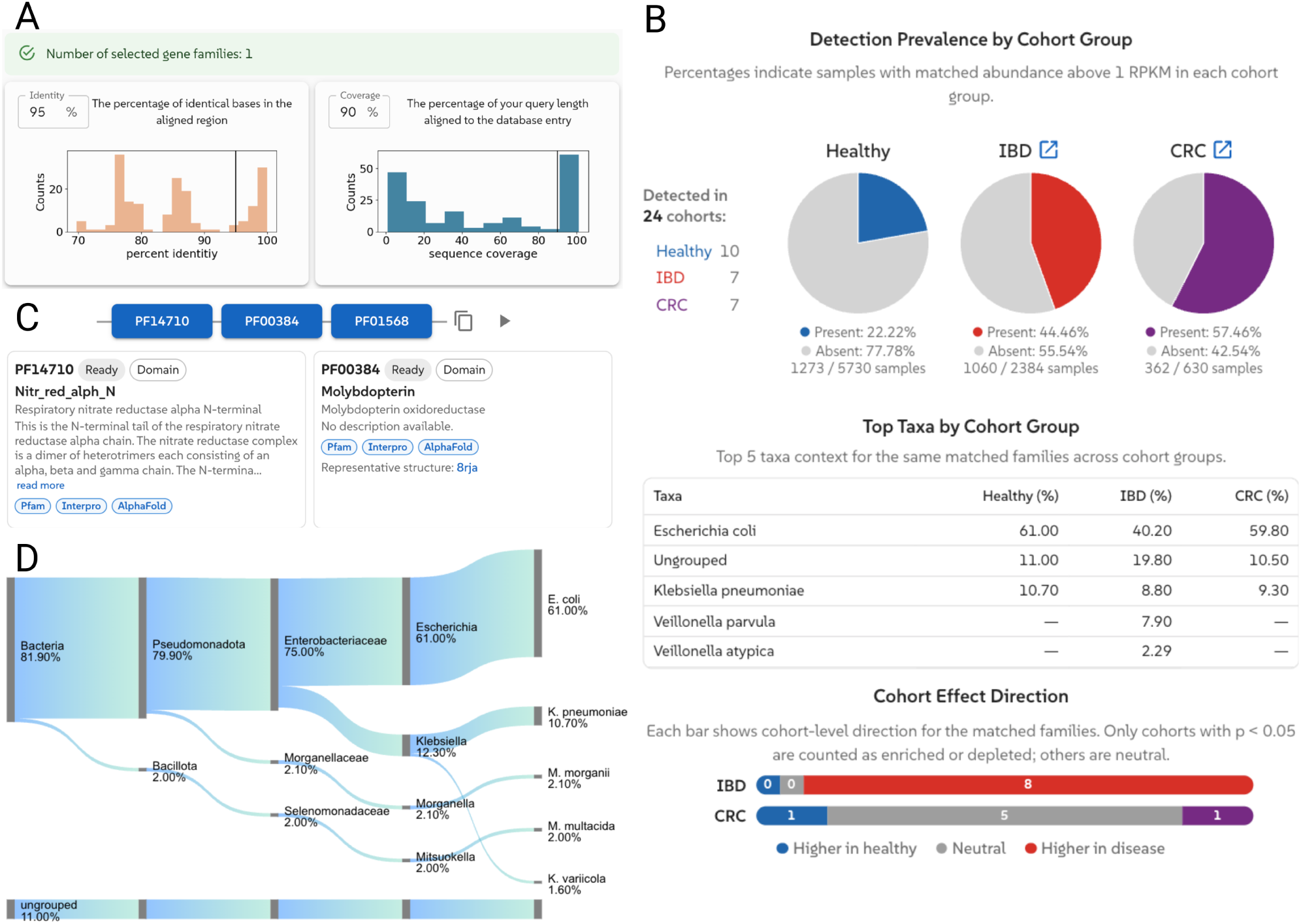
MetaGEAR Explorer interface: *narG* search results and overview. (**A**) BLAST alignment statistics for the *narG* sequence search (percent identity, sequence coverage, E-value, bit score) with adjustable thresholds. (**B**) Cross-cohort disease association for the NarG domain search (160 gene families): detection prevalence by cohort group, top taxa, and cohort effect direction bars. (**C**) Pfam domain architecture of matched families, cross-referenced with InterPro and linked to AlphaFold structures. (**D**) Taxonomic composition (Healthy) as a Sankey diagram (consensus GTDB-Tk/MetaPhlAn 3 taxonomy). Data plots derived from these views appear in Figure 2.

**Supplementary Figure 3.**
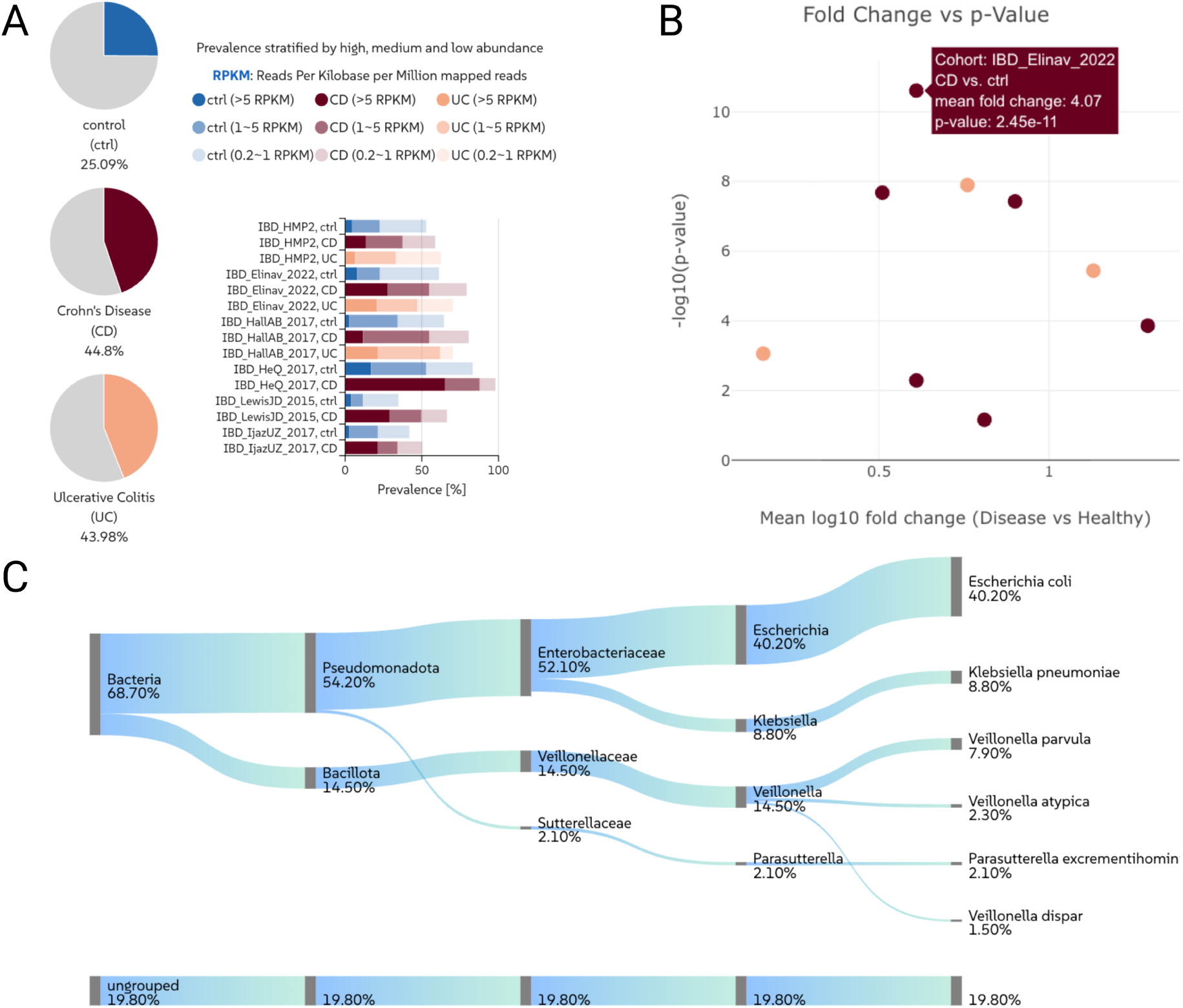
MetaGEAR Explorer interface: *narG* disease details (IBD). (**A**) Sub-group prevalence (non-IBD Control, CD, UC) with per-cohort bars stratified by abundance level. (**B**) Disease-association scatter (mean fold change vs. −log_10_ p) per cohort comparison. (**C**) IBD-filtered taxonomic composition as a Sankey diagram. Data plots derived from these views appear in Figures 2 and 3.

**Supplementary Figure 4.**
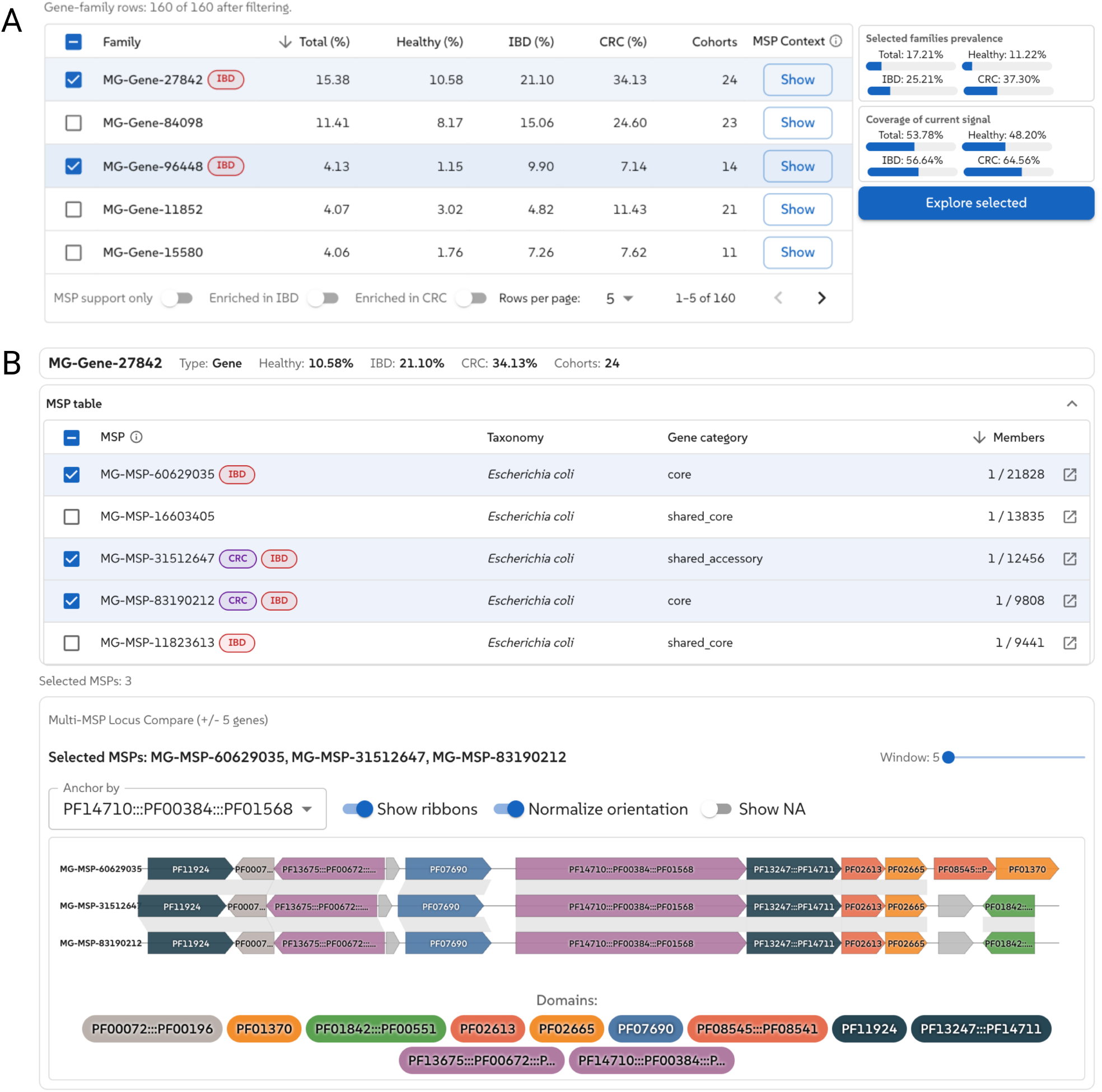
MetaGEAR Explorer interface: gene/protein family explorer. (**A**) Matched-family table with per-group prevalence, cohort count, and MSP context; disease-enriched families are labelled. (**B**) MSP context for a selected family and a multi-MSP locus comparison aligning genomic neighborhoods of three *E. coli* MSPs, anchored on the NarG domain architecture with ribbon connectors. Data plots derived from these views appear in Figure 3.

**Supplementary Figure 5.**
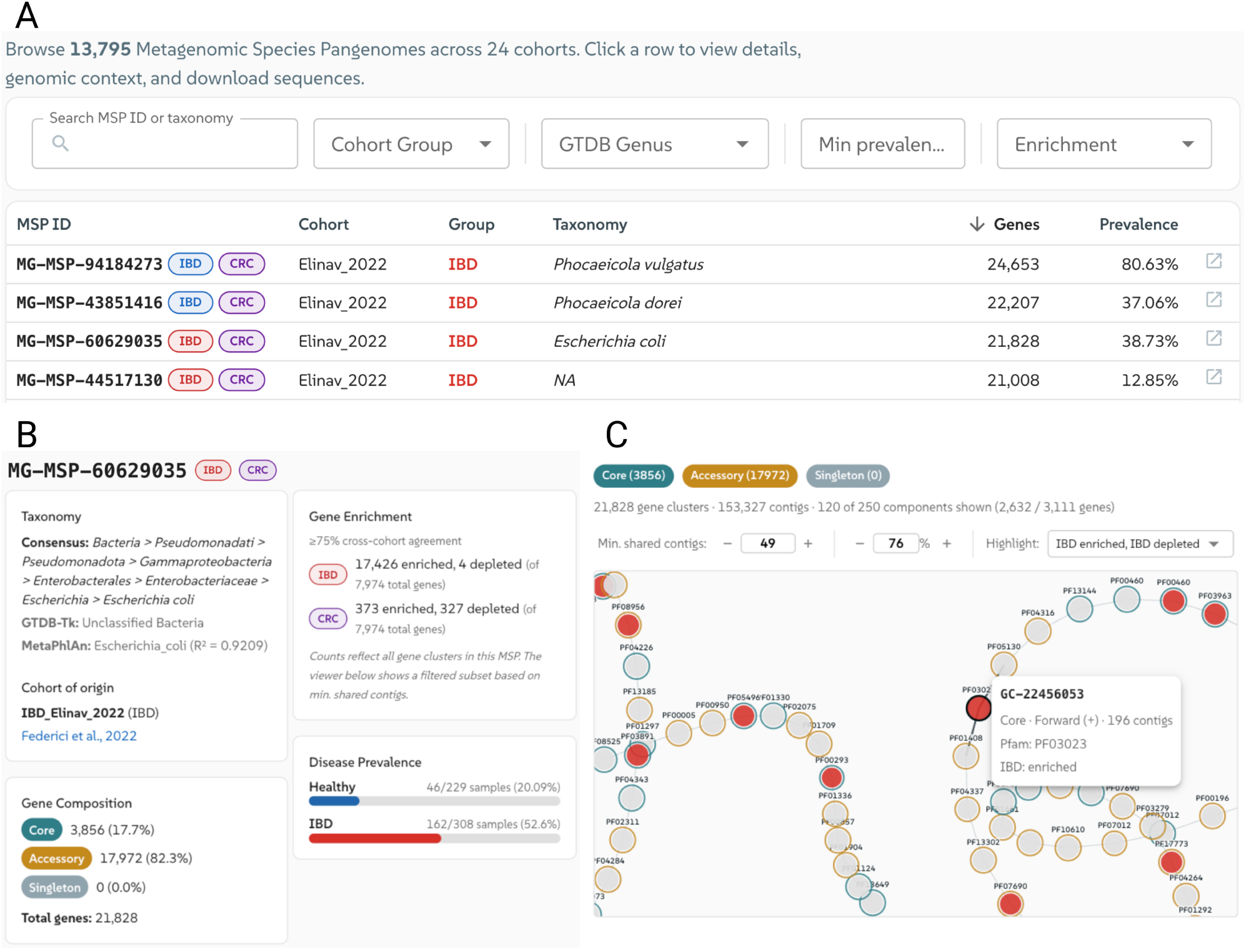
MetaGEAR Explorer interface: MSP catalog browser and pangenome viewer. (**A**) MSP catalog browser (13,795 MSPs) with consensus taxonomy and sortable, filterable columns. (**B**) MSP detail page for an *E. coli* MSP (MG-MSP-60629035): taxonomies, gene composition, and disease prevalence. (**C**) MSP Viewer pangenome graph (force-directed); node fill indicates disease association and rings denote gene category. Data plots derived from these views appear in Figure 3.

**Supplementary Figure 6.**
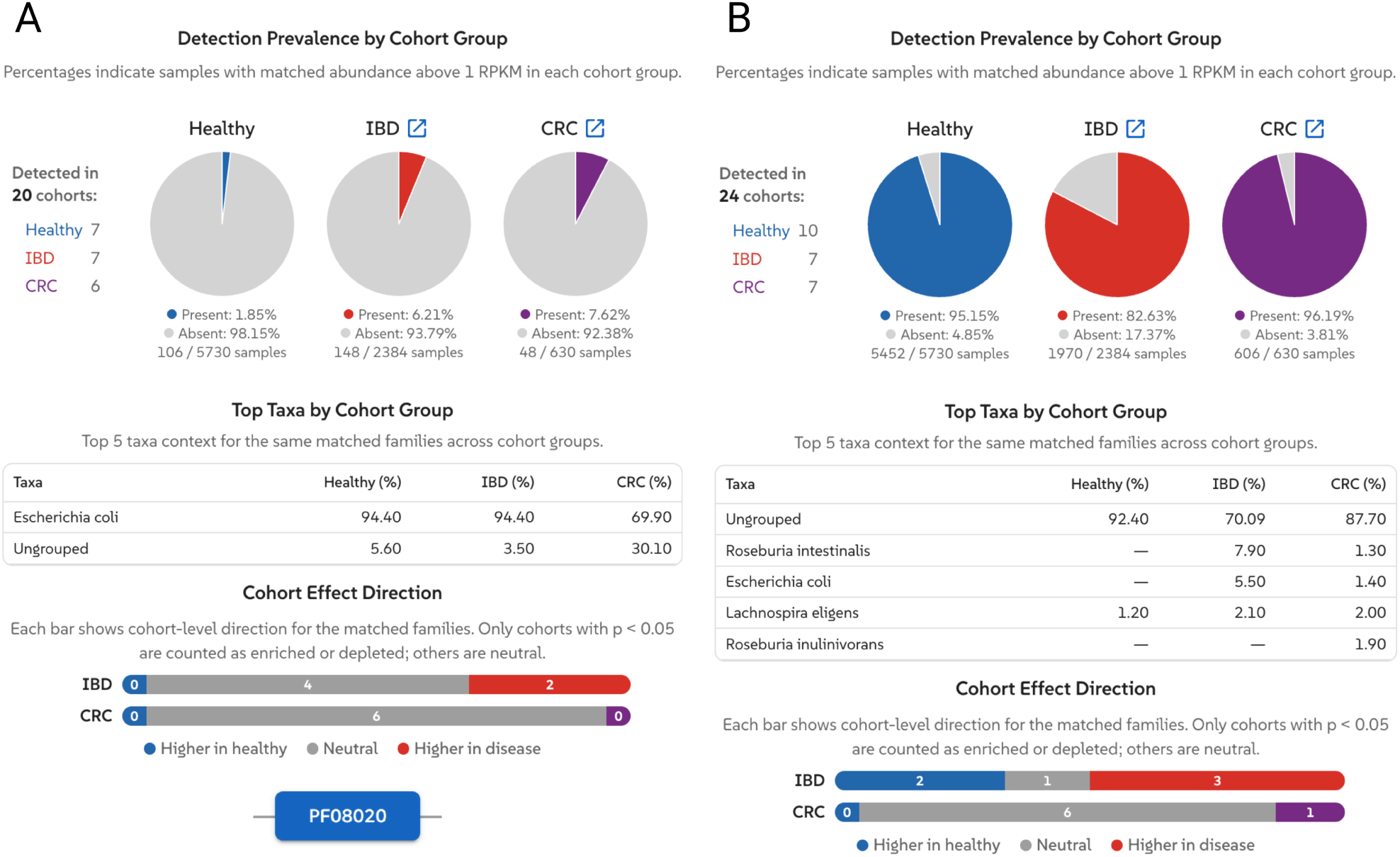
MetaGEAR Explorer interface: web-based discovery of colibactin immunity genes. (**A**) Cross-cohort disease association from a BLASTp search of two canonical ClbS sequences (1.85% Healthy, 6.21% IBD, 7.62% CRC; *E. coli*-dominated). (**B**) Cross-cohort disease association from the PF08020 (DUF1706) domain search (1,121 gene families; 95.15% Healthy, depleted to 82.63% in IBD). Data plots derived from these views appear in Figure 4.

**Supplementary Figure 7.**
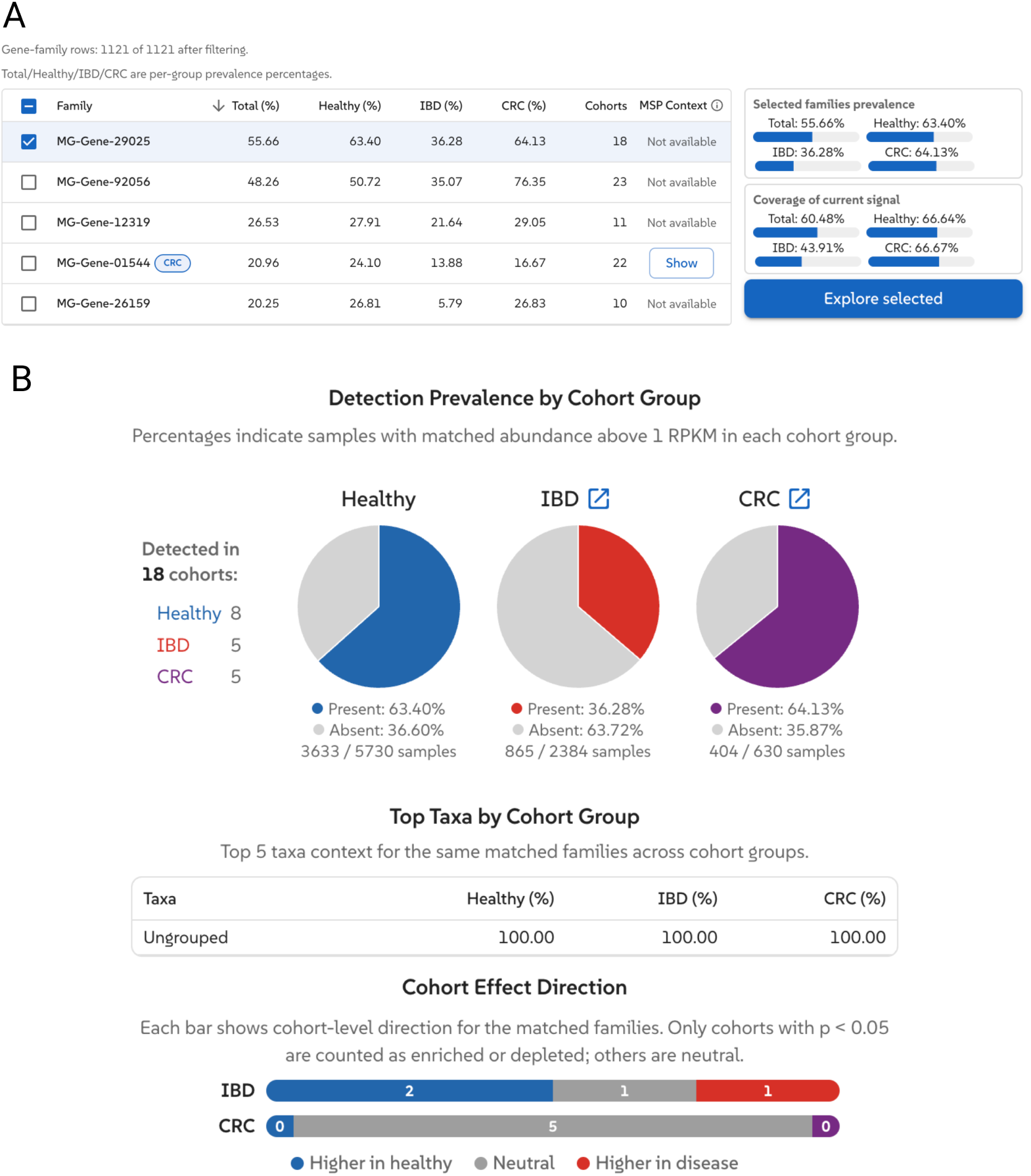
Dominant uncharacterized DUF1706 gene family (MG-Gene-29025) from the PF08020 domain search. (**A**) Matched gene family table from the PF08020 domain search (1,121 gene families), sorted by total prevalence. MG-Gene-29025 is the most prevalent gene family (55.66% total), detected in 18 of 24 cohorts. Its MSP context is unavailable, indicating no metagenomic species pangenome assignment. Selection coverage shows this single gene family accounts for ∼60% of the total PF08020 signal. (**B**) Disease-association profile for MG-Gene-29025: prevalence is 63.40% in healthy controls and 64.13% in CRC, but drops to 36.28% in IBD. Taxonomy is 100% ungrouped across all cohort groups, meaning the carrier organism(s) cannot be assigned to any species in the current catalog. The cohort effect direction for IBD shows 2 cohorts with higher prevalence in healthy versus 1 in disease, consistent with the prevalence drop observed in the pie charts.

## Appendix C Test Input Data

### C.1 NarG: Nucleotide sequence

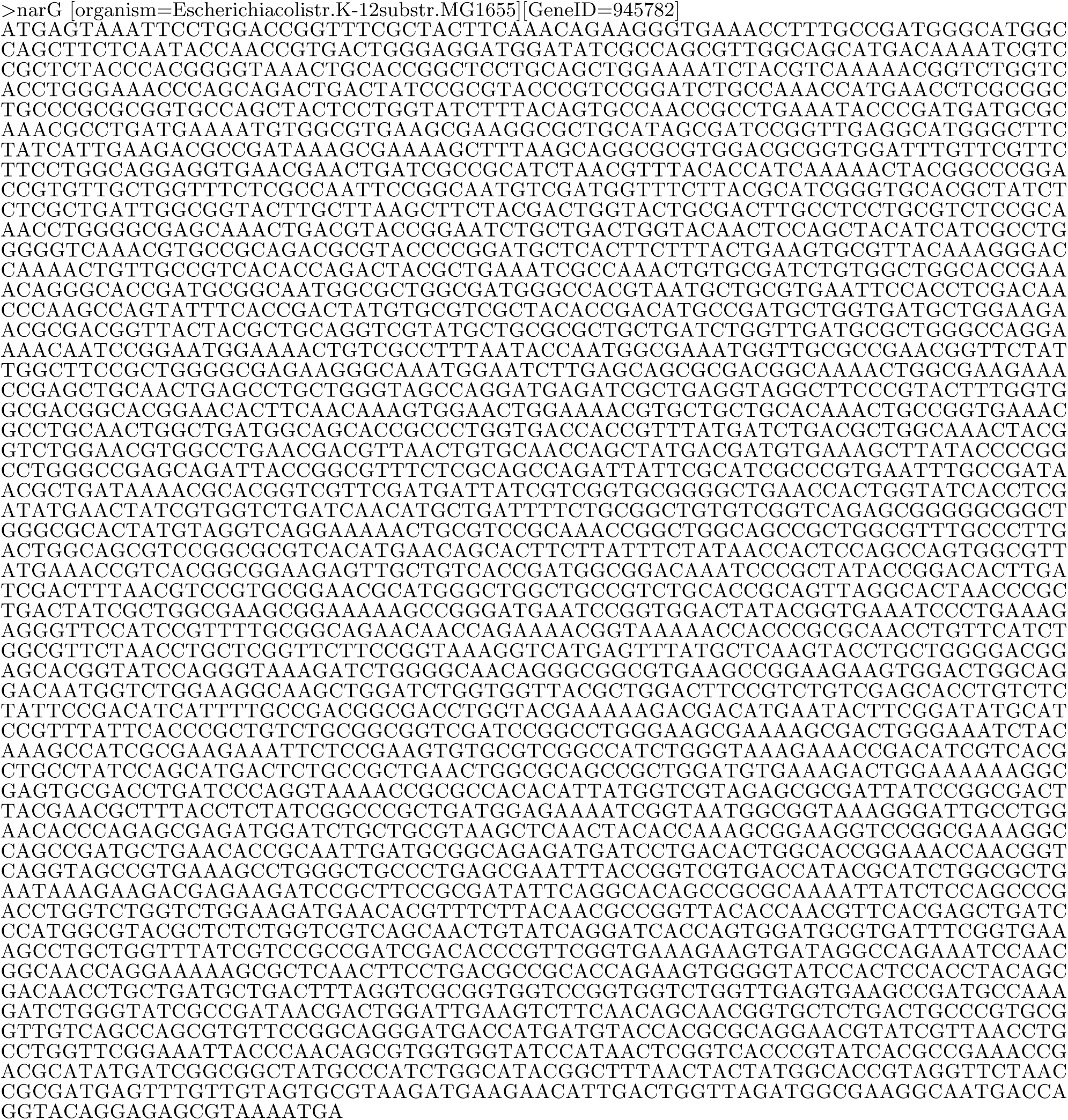

### C.2 NarG: Amino acid sequence

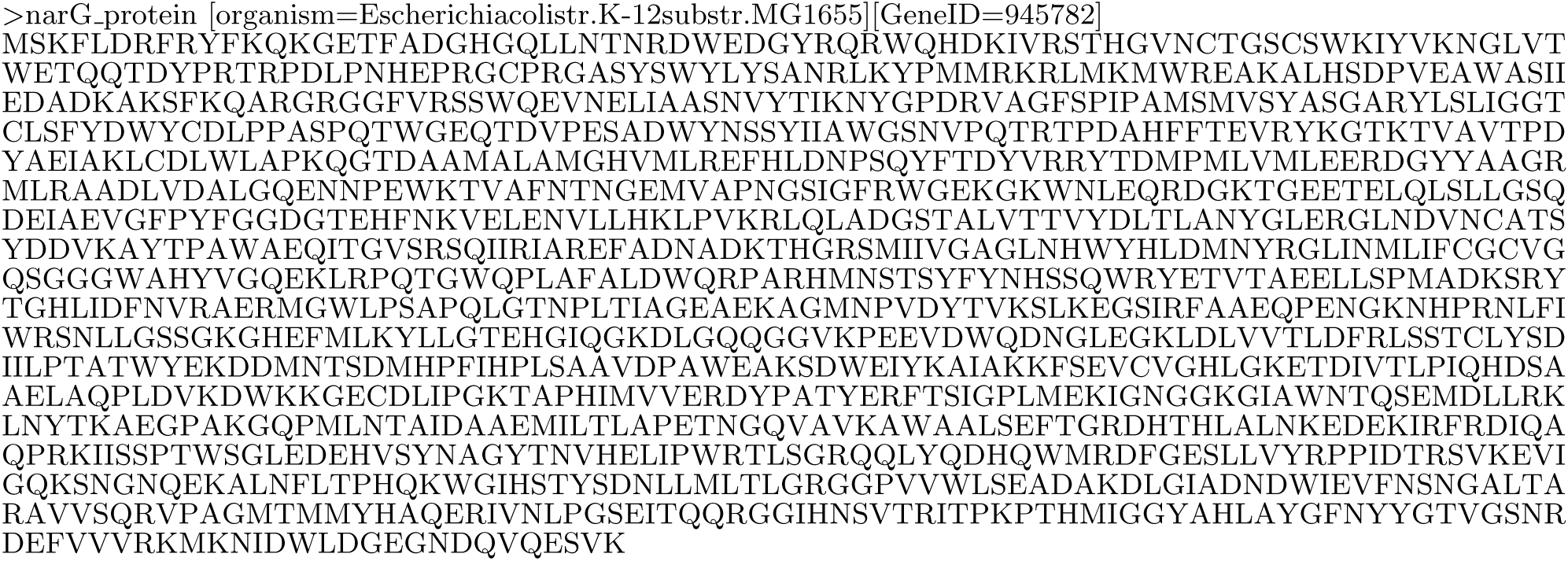

### C.3 NarG-like: Functional domain configuration

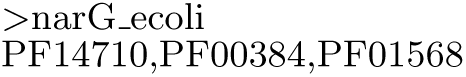

### C.4 ClbS: Amino acid sequences (colibactin self-protection protein)

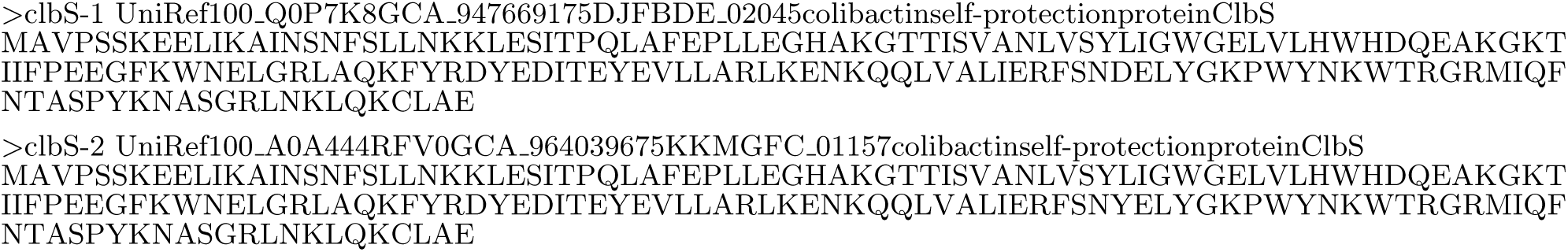

### C.5 ClbS / DUF1706: Functional domain configuration

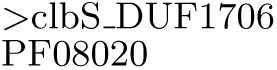

